# A PACAP-activated network for secretion requires coordination of Ca^2+^ influx and Ca^2+^ mobilization

**DOI:** 10.1101/2024.01.03.574069

**Authors:** Xiaohuan Chen, Nicole A. Bell, Breanna L. Coffman, Agustin A. Rabino, Rafael Garcia-Mata, Paul J. Kammermeier, David I. Yule, Daniel Axelrod, Alan V. Smrcka, David R. Giovannucci, Arun Anantharam

## Abstract

Chromaffin cells of the adrenal medulla transduce sympathetic nerve activity into stress hormone secretion. The two neurotransmitters principally responsible for coupling cell stimulation to secretion are acetylcholine and pituitary adenylate activating polypeptide (PACAP). In contrast to acetylcholine, PACAP evokes a persistent secretory response from chromaffin cells. However, the mechanisms by which PACAP acts are poorly understood. Here, it is shown that PACAP induces sustained increases in cytosolic Ca^2+^ which are disrupted when Ca^2+^ influx through L-type channels is blocked or internal Ca^2+^ stores are depleted. PACAP liberates stored Ca^2+^ via inositol trisphosphate receptors (IP3Rs) on the endoplasmic reticulum (ER), thereby functionally coupling Ca^2+^ mobilization to Ca^2+^ influx and supporting Ca^2+^-induced Ca^2+^-release. These Ca^2+^ influx and mobilization pathways are unified by an absolute dependence on phospholipase C epsilon (PLCε) activity. Thus, the persistent secretory response that is a defining feature of PACAP activity, *in situ*, is regulated by a signaling network that promotes sustained elevations in intracellular Ca^2+^ through multiple pathways.

## Introduction

The adrenal medulla is a core effector of the sympathetic nervous system in the periphery (Cannon, 1940; Carmichael and Winkler, 1985). In response to activation by sympathetic splanchnic nerves, the medulla releases a cocktail of hormones into the bloodstream for circulation throughout the body. Chromaffin cells are the secretory units of the adrenal medulla (Carmichael and Winkler, 1985). Chromaffin cells synthesize, store, and secrete epinephrine, norepinephrine, and various bioactive peptides that modulate cardiovascular tone, increase glucose availability, reorient the flow of blood, and control digestion (Cannon, 1940; Goldstein, 2010). In addition to their well-defined role in the sympathetic stress response, chromaffin cells have served as a veritable “Rosetta Stone” for our understanding of the basic rules of exocytosis in other systems. It is on account of the chromaffin cell that the basic features of exocytosis, including its Ca^2+^ dependence, its regulation by phospholipids and proteins, and the properties of its fusion pores, are now so well understood (Anantharam and Kreutzberger, 2019). Similar features have subsequently been identified in other secretory cell types, from neurons in the brain to beta cells in the pancreas (Hastoy et al., 2017; Neher, 2006).

Chromaffin cell secretion is primarily controlled by two neurotransmitters – acetylcholine (ACh) and pituitary adenylate cyclase activating polypeptide (PACAP) (Carbone et al., 2019; Guerineau, 2019). While the actions of acetylcholine in the medulla have been interrogated for many decades, beginning with the work of Feldberg in the 1930s (Feldberg et al., 1934), the studies of PACAP-stimulated secretion are still very much at a nascent stage. Indeed, our knowledge of PACAP and its physiological effects, particularly in the context of stress transduction, are much more advanced elsewhere in the body than it is in the adrenal (Rajbhandari et al., 2023; Riser and Norrholm, 2022). In the central nervous system (CNS), PACAP was originally identified as a peptide in the hypothalamus and shown to stimulate adenylate cyclase activity in the anterior pituitary gland (Miyata et al., 1989). Subsequent studies established PACAP as a powerful regulator of the hypothalamic-pituitary-adrenal (HPA) circuit, with its expression being necessary for the release of adrenocorticotropic hormone, corticosterone, and corticotropic-releasing hormone (Riser and Norrholm, 2022). Recent work has revealed a strong connection between enhanced PACAP secretion, single nucleotide polymorphisms in the PACAP receptor, and post-traumatic stress disorder in human populations (Ressler et al., 2011; Stevens et al., 2014; Wang et al., 2021).

In the adrenomedullary system, it is now firmly established that PACAP is released from splanchnic nerve terminals onto chromaffin cells, causing sustained epinephrine secretion even in the background of nicotinic receptor desensitization (Hill et al., 2011; Stroth et al., 2013). In the periphery, as in the CNS, the secretory function of PACAP is critical for physiological adaptations to stress (Guerineau, 2019; Hamelink et al., 2002). Mice harboring a targeted deletion of the PACAP gene, and subjected to a strong sympathetic stressor, exhibit substantially reduced epinephrine secretion compared to WT controls (Hamelink et al., 2002).

In contrast to acetylcholine – which causes membrane depolarization, the opening of voltage-gated channels, and dense core vesicle fusion – the mechanisms coupling PACAP stimulation in the chromaffin cell to fusion remain poorly understood (Carbone et al., 2019; Eiden et al., 2018). The specific intracellular consequences of PACAP stimulation appear to vary by cell-type and system, with Gα_q_ and Gα_s_ proteins being implicated in its signaling pathways (Rajbhandari et al., 2023). However, it has been clear from the earliest studies of PACAP action that the secretion it provokes is Ca^2+^-dependent (Chowdhury et al., 1994; Przywara et al., 1996; Watanabe et al., 1992). While much of the recent attention has been focused on the role of low voltage-activated T-type channels, it is unclear whether this is the only, or even the predominant, pathway linking Ca^2+^ to PACAP-evoked secretion (Hill et al., 2011; Kuri et al., 2009).

In this study, the mechanisms by which PACAP elevates intracellular Ca^2+^ and triggers exocytosis in chromaffin cells were systematically investigated. This was done with a combination of high-resolution imaging approaches, sensitive optical reporters of cellular activity, and pharmacological and genetic perturbations of proteins implicated in the PACAP signaling pathway. The experiments demonstrate that PACAP-evoked secretion requires the liberation of Ca^2+^ from the endoplasmic reticulum (ER) as well as Ca^2+^ influx from the extracellular space through L-type channels. Pharmacological inhibition of IP3 receptors, or shRNA-based knockdown of the IP3R1 isoform, largely eliminated the Ca^2+^ signals normally stimulated by PACAP. PACAP failed to elevate Ca^2+^ or evoke secretion in cells that lacked PLCε – an enzyme that produces IP3. These data support a model where PACAP, signaling through PLCε, elevates cytosolic Ca^2+^ and promotes IP3R1-mediated Ca^2+^-induced Ca^2+^ release (CICR) (Lock et al., 2018; Lock and Parker, 2020; Marchant and Parker, 2001). Thus, the persistent secretory response that is a defining feature of PACAP activity, *in situ*, is regulated by a cellular signaling network that causes sustained elevations in intracellular Ca^2+^.

## Materials and Methods

### Animals

C57BL/6J mice (referred to as “wild-type” or WT) were obtained from Jackson Labs (Bar Harbor, ME). PLCε -/-mice (also referred to as “KO”) were generated by Smrcka and colleagues (Atchison et al., 2020; Wang et al., 2005). The mice were maintained in group housing with 2-5 mice per ventilated cage under a 12-hour dark/12-hour light cycle with full access to food and water. All animal procedures and experiments were conducted following the stipulated University of Toledo (400138) IACUC protocol.

### Mouse chromaffin cell preparation

Mouse chromaffin cells were isolated and cultured following previously established protocols (Morales et al., 2023). In brief, 2–4-month-old male or female mice were anesthetized using isoflurane and euthanized via cervical dislocation. Adrenal glands were extracted and transferred to dishes containing ice-cold mouse buffer (148 mM NaCl, 2.57 mM KCl, 2.2 mM K_2_HPO_4_•3H_2_O, 6.5 mM KH_2_PO_4_, 10 mM glucose, 5 mM HEPES free acid, 14.2 mM mannitol, adjust pH to 7.2). After careful removal of the cortex, the medullae were rinsed three times in 100 μL drops of papain enzyme solution (450 units/ml Papain; Worthington Biochemical, Cat# LS003126), 250 μg/ml bovine serum albumin (BSA; Sigma-Aldrich, Cat# A7906), and 75 μg/ml dithiothreitol (Roche, Cat# 10708984001) in mouse buffer) and then transferred to a 15-mL falcon tube containing 0.5 mL of papain solution placed in a water bath shaker for 15 minutes at 37°C shaking at 140 rpm. After 15 minutes, the digesting solution was mostly removed and replaced by 0.5 mL of collagenase enzyme solution (250 μg/ml BSA, 3.75 mg/ml collagenase (Sigma-Aldrich, Cat# C0130), and 0.15 mg/mL DNase I (Sigma-Aldrich, Cat# DN25) in mouse buffer). Digestion was continued for another 15 minutes at 37°C shaking at 140 rpm. Post-digestion, the medullae were transferred to antibiotic-free culture medium (Dulbecco’s Modified Eagle’s Medium/F12 (DMEM/F12; ThermoFisher Scientific, Cat# 11330-032) supplemented with 10% Fetal Bovine Serum (FBS) (ThermoFisher Scientific, Cat# 11403-028), triturated using a pipette, and centrifuged at 300g for 2.5 min. The resultant pellet (from 2-4 glands) was resuspended by 300 μL of antibiotic-free medium, and the mixed cell suspension was equally divided into two coated dishes. Before the start of the cell preparation, 35 mm glass bottom dishes (10 mm glass diameter; MatTek, Cat# P35G-1.5-10-C) were pre-coated with Matrigel (Corning, Cat# 356230) diluted in DMEM/F12 (1:7) for 1-1.5 hours after which the dishes were washed with DMEM/F12. The cells were cultured in an incubator (37°C, 5% CO_2_) for approximately 4 hours. A culture medium with antibiotics was then added to a final volume of 2 mL (DMEM/F12 supplemented with 10% FBS, 9.52 unit/ml Penicillin, 9.52 µg/ml Streptomycin, and 238 μg/ml Gentamicin (ThermoFisher Scientific, Cat# 15140-122 and Cat# 15710-064)). The media was replaced the day after plating and experiments were conducted between 18 and 48 hours after plating.

### Plasmids, Transfection, Fluorescent dyes

The GCaMP5G plasmid was procured from Addgene (Cat# 31788). Lck was fused to GCaMP5G as described in Shigetomi et al. (2010). The Neuropeptide Y (NPY) pHluorin was generously provided by Dr. Ronald W. Holz (University of Michigan, Ann Arbor, MI). The ER-GCaMP6-150 plasmid was provided by Dr. Timothy Ryan (Weill Cornell, New York, NY). The PLCε-FLAG-P2A-mCherry plasmid was synthesized and subsequently cloned into a pCMV-script EX vector by GenScript. For rescue experiments involving PLCε KO chromaffin cells, cells exhibiting overexpression of PLCε-FLAG were identified based on their mCherry fluorescence. The scrambled shRNA and IP3R1 shRNA plasmids were purchased from OriGene; the 29mer sequence of scrambled shRNA (Cat# TR30015) is 5’-GCACTACCAGAGCTAACTCAGATAGTACT-3’, the 29mer sequence of IP3R1 (also called Itpr1) shRNA (Cat# TF517036) is 5’-TCAGCACCTTAGGCTTGGTTG-ATGACCGT-3’, and the transfected cells were identified based on their red fluorescence encoded by shRNA vectors. For transfections, cell pellets were resuspended in 110 μL of sucrose-based buffer (250 mM sucrose and 1 mM MgCl_2_ in DPBS (Gibco, Cat# 10010-023)). The desired plasmid was added to the mixture (1.5 μg/gland). The suspended cells were transiently transfected by electroporation with a single pulse (1050 mV, 40 ms) using the neon transfection system (Invitrogen, Cat# MPK5000 and Cat# MPK10096). After transfection, the mixture was mixed with 200 μL antibiotic-free culture medium and divided into 2 Matrigel-coated dishes. After approximately 4 hours, 1.5 ml medium with antibiotics was added to the cell.

In some cases, cells were not transfected but incubated with fluorescent dyes for imaging Ca^2+^, the cell membrane, or vesicle fusion. For cell membrane staining, cells were incubated with 5 μg/ml CellMask™ (Thermo Fisher, Cat# C10046) for 10 min, and subsequently washed with PSS 3 times before imaging. For imaging of Ca^2+^ puffs, cells were incubated with 1 µM of the membrane-permeant fluorescent Ca^2+^ indicator Cal-520/AM (AAT Bioquest, Cat# 21130) in normal PSS for 30 min. For labeling vesicles, cells were incubated with 10 μM FFN511 (Abcam, Cat# ab120331) in PSS buffer for 30 min and washed with PSS prior to imaging on the TIRF microscope and offline analysis.

### Western blotting

Adrenal glands were carefully dissected and transferred to dishes containing ice-cold mouse buffer. The adrenal cortex was surgically trimmed to isolate the adrenal medulla. Medulla were then combined and lysed in an Eppendorf tube containing 150 µl MPER buffer (Thermo Scientific, Cat# 78501) with a protease inhibitor cocktail (Thermo Scientific, Cat# 78444). The lysing process took place on ice for 30 minutes, with intermittent vortexing every 5 minutes. The lysed samples were centrifuged at 12,000 × g for 15 min at 4°C, with the supernatant transferred to a fresh tube for protein concentration measurement via BCA assay (Thermo scientific, Cat# A53227). Each sample, containing 20 µg of protein, was loaded onto a gradient (4–12%) Bolt Bis-Tris plus gel (Thermo scientific, Cat# NW04122BOX) for separation in 1X Bolt MES SDS running buffer (Thermo scientific, Cat# B0002) for 2 h at 120 V. Proteins were then transferred to a PVDF membrane in 1X Bolt transfer buffer (Thermo scientific, Cat# BT0006) for overnight at 25V. Membrane was blocked with 5% non-fat milk in TBST for 1 h at room temperature. Primary antibodies were added and incubated overnight at 4°C. Membrane was washed 3X with TBST for 10 min each and HRP-conjugated secondary antibodies were added and incubated at room temperature for 1 h. Membrane was washed again with 3X with TBST for 10 min each. Blots were then developed using an ECL reagent (Thermo scientific, Cat# 34580) in a box western blot imager (G: Box chemi, Syngene). Antibody information is as follows: rabbit anti-IP3R1, 1:5,000 (Yule lab in-house, clone CT1), mouse anti-beta actin, 1:2000 (Thermo scientific, Cat# MA1-140, RRID: AB_2536844, clone 15G5A11/E2), goat anti-rabbit IgG (H+L) secondary antibody, HRP, 1:2000 (Thermo scientific, Cat# 31460, RRID: AB_228341), goat anti-mouse IgG1 cross-adsorbed secondary antibody, HRP, 1:2000 (Thermo scientific, Cat# A10551, RRID: AB_2534048). For the IP3R1 knock down experiment, cells from 10 glands were collected together and transfected with either 10 μg scrambled shRNA or 10 μg IP3R1 shRNA plasmid. Cells were then seeded on 12-well plates. After 24 hours, complete media was removed, and cells were cultured with new complete media containing 100 ug/ml puromycin (Sigma, Cat# P8833) for an additional 3 days to kill any untransfected cells. Cells were then lysed and subjected to western blot analysis as described above.

### Real-time PCR

After rinsing the cultured chromaffin cells with PBS thrice, the cells were lysed with 300 µl TRIzol reagent (Thermo scientific, Cat# 15596026). Following the manufacturer’s guidelines, total RNA was extracted and 200 ng of total RNA per sample was converted into cDNA using the SuperScript IV VILO Master Mix kit (Thermo scientific, Cat# 11756500). Quantitative PCR was run with PowerUp SYBR green Master Mix (Thermo scientific, Cat# A25742) on a Quant Studio 3 detection system (Applied Biosystems, Waltham, MA, USA). The 2^−ΔCT^ method was used to quantify relative mRNA expression levels which were normalized to beta-actin. The assays were performed in both biological and technical triplicate. Custom primers were prepared by Integrated DNA Technologies, the sequences of which are specified in Table 1 below.

**Table 1.**
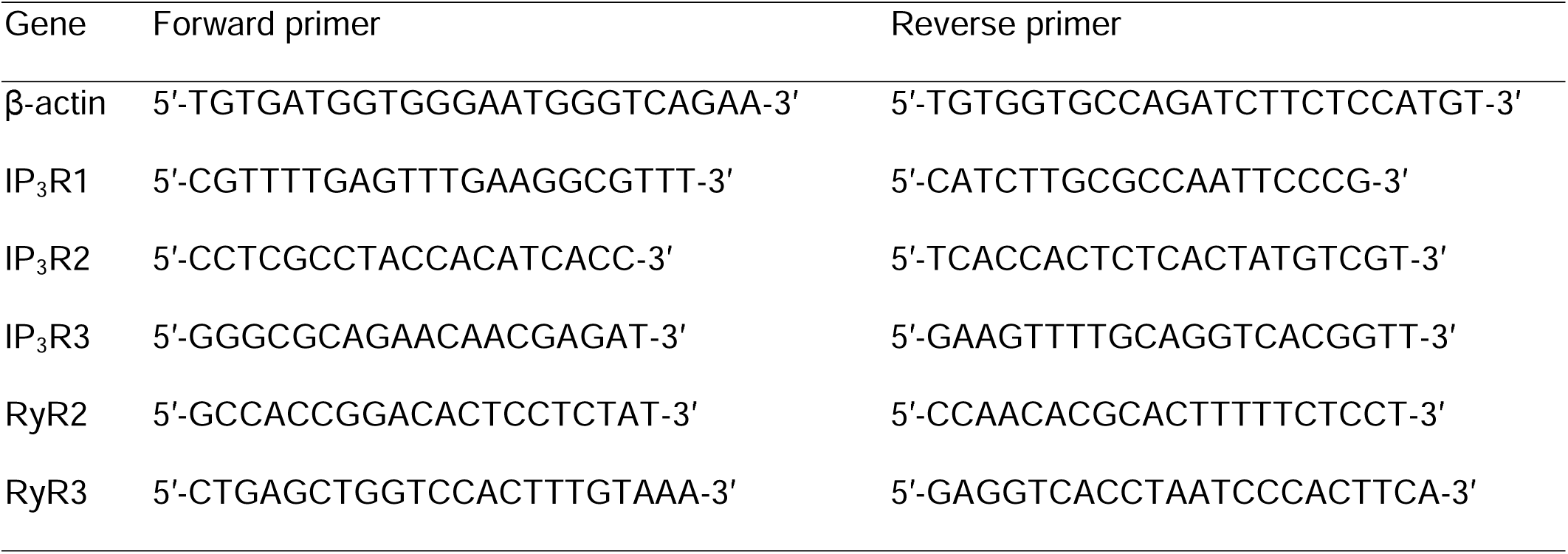

### TIRF Microscopy

TIRF imaging was performed utilizing an Olympus cellTIRF microscope system (Olympus, USA), configured for two-line (488nm/561nm) operation. The microscope was equipped with high numerical aperture (NA 1.49) TIRF oil-immersion objectives (60x and 100x) and was sometimes used with an additional 2x lens in the emission path situated between the microscope and the camera (Andor Technology, iXon Ultra 897). The resultant pixel size in the images was between 80 – 160 nm.

All TIRF experiments were executed at a controlled temperature range of 35-37°C, achieved by placing the sample dish on a temperature controller platform (Warner instruments, Cat# TC-324C). The physiological salt solution (PSS) buffer, composed of 145 mM NaCl, 5.6 mM KCl, 2.2 mM CaCl_2_, 0.5 mM MgCl_2_, 5.6 mM glucose, and 15 mM HEPES (pH 7.4), was preheated to 37°C prior to experimentation. The Ca^2+^-free PSS buffer was also supplemented with 300 uM EGTA (EMD Millipore, Cat# 324626). The culture medium was replaced with PSS immediately before the experiment. The replacement of the PSS bath was accomplished through a gravity-based system, functioning at a fluid flow rate of 3-5 mL per minute. Chromaffin cells were individually stimulated utilizing a 100-μm inner diameter needle (ALA Scientific Instruments, Cat# ALA QTP) in conjunction with a positive-pressure perfusion system (ALA Scientific Instruments, ALA-VM4).

### Confocal microscopy

All confocal images were taken using the Leica Stellaris 5 laser scanning confocal equipped with an HC PL APO 63×/1.40 OIL CS2 objective and LAS X software. An additional 6x zoom-in was applied to the field of interest using the microscope’s software. All images were captured in sequential scan mode according to the laser channels, with excitation provided by 405-, 488-, 594-, and 647 nm lasers. All images were further processed using ImageJ Fiji software.

### Live Cell cAMP, R-GECO, and DAG Assay

Cells were transduced with either green cADDis (cAMP) upward sensor (Montana Molecular, Cat# U0200G), red GECO Ca^2+^ upward sensor (Montana Molecular, Cat# U-0600R), or red DAG downward sensor (Montana Molecular, Cat# D-0300R) following the manufacturer’s protocol. Briefly, cells were infected with 15 μL of BacMam sensor stock in 150 μl of media, supplemented with 2 mM Sodium Butyrate (Montana Molecular), for 30 minutes at room temperature, away from light. This was followed by a 4 to 6-hour incubation period in a 37°C tissue culture incubator. The BacMam solution was then removed and replaced with media containing 2 mM Sodium Butyrate for a duration of 18 to 48 hours. Imaging was performed on the TIRF microscope with images acquired every 5 seconds for a duration of 5 minutes (100 ms exposure time) with PACAP perfused after the initial 20-25 seconds.

### Imaging protocols for pharmacology

During experiments requiring sequential application of pituitary adenylate cyclase-activating polypeptide 1-38 (PACAP, synthesized at University of Iowa by J. Galpin and C. Ahern (Chen et al., 2023)) and calcium channel blockers, cells underwent an initial 2.5 s exposure to PSS, followed by a 30 s stimulation with PACAP. Subsequently, cells were exposed to a combination of PACAP and lower concentrations of blockers (as indicated in figures) for 30 s, then PACAP and higher concentrations of blockers, followed by a 30 s wash with PSS, and finally another 30 s stimulation with PACAP. The L-type calcium channel blocker, nifedipine (Cat# N7634), and the N-type calcium channel blocker, ω-Conotoxin GVIA (Cat# C9915), were purchased from Sigma. The P/Q-type calcium channel blocker, ω-Agatoxin TK (Cat# STA-530), was purchased from Alomone Labs.

In experiments which utilized caffeine (Alomone Labs, Cat# C-395), cells were initially perfused with PSS for 2.5 s, then perfused with 40 mM caffeine for 3 min, followed by PACAP for 30 s. Thereafter, cells were perfused with PSS for an additional 5 min, and finally subjected to another 30 s of PACAP perfusion. In all other experiments utilizing inhibitors, including Xestospongin C (Xesto C, Abcam, Cat# ab120914), Cyclopiazonic acid (CPA, Sigma, Cat# C1530), and ryanodine (Tocris, Cat# 1329), cells were incubated with inhibitors for a minimum of 1 minute prior to stimulation with either 500 nM PACAP or 1 µM DMPP (Sigma, Cat# D5891).

The experiments involving ER-GCaMP6-150 were performed as follows. First, the cell was perfused with normal PSS containing 2.2 mM Ca^2+^ for 1 min, followed by perfusion with 500 nM PACAP in normal PSS for 1 min. Next, the cell was perfused for 1 min with PACAP without Ca^2+^ followed by perfusion with normal PSS for 1 min. Finally, the cell was perfused with 40 mM caffeine for 20 s.

### Immunocytochemistry

Freshly dissociated chromaffin cells were cultured in MatTek dishes for a duration of 24 to 48 hours after which the culture medium was removed, and the cells were washed three times with PBS (Gibco, Cat# 10010023). Following the washing process, cells were fixed with 4% PFA for 10 minutes at room temperature. They were then washed three more times with PBS and blocked for 1 hour at room temperature using 10% BSA (Sigma, Cat# A7906) in 1X PBST (PBS + 0.2% Triton X-100, Sigma, Cat# T8787). The cells were then incubated with primary antibodies. These included Rabbit anti-IP3R1 at 1:1000 (a gift from Dr. David Yule, clone CT1), Rabbit anti-IP3R1 at 1:200 (Thermo Fisher, Cat# PA1-901, RRID:AB_2129984), and Mouse anti-KDEL at 1:500 (Enzo Life Sciences, Cat# ADI-SPA-827, RRID:AB_10618036, clone 10C3). The antibodies were diluted in 2% BSA in 1X PBST and the cells were incubated with them overnight at 4 °C. The following day, the cells were washed three times with PBST for 15 minutes each. Subsequently, they were incubated with Alexa Fluor 488-donkey anti-rabbit IgG (H + L) at 1:500 (Thermo Fisher Scientific, Cat# A-21206, RRID:AB_2535792) and Alexa Fluor 568-donkey anti-mouse IgG (H + L) at 1:500 (Thermo Fisher Scientific, Cat# A10037, RRID:AB_2534013) in 2% BSA in 1X PBST for one hour at room temperature. Following the incubation, the cells were washed three times with PBST for 15 minutes each and mounted with DAPI Fluoromount G (SouthernBiotech, Cat# 0100-20) for imaging.

### Colocalization Analysis

The JACoP plugin on ImageJ was used to compute the Manders coefficients of the green (Alexa 488) and red (Alexa 594) areas of interest. These areas corresponded to the antibody labeled IP3R1 and KDEL proteins, respectively. Each cell was then exposed to a thresholding setting using the JACoP ImageJ plugin, as detailed by Bolte and Cordelieres (2006). The plugin automatically calculated the M1 and M2 overlap coefficients.

### Imaging and image analysis of NPY-pHluorin and FFN511 fusion events

Fusion events were visually identified. Regions of Interest (ROIs) with a radius of 240 nm were then drawn using the Time Series Analyzer v3.0 plugin on Fiji software. Image acquisition was performed at a frequency of 50 frames per second. The duration of NPY-pHluorin and FFN511 release was assessed with a custom-written program in Interactive Data Language (IDL; ITT, Broomfield, CO)(Morales et al., 2023).

### Calcium Imaging and Analysis

#### Analysis of global calcium signals (amplitude, total duration, and total area under the curve)

Global Ca^2+^ signals were measured in cells expressing Lck-GCaMP5G or Cal-520. The frame rate for these experiments was typically 20 frames per second. The data preparation, ROI selection, and analyses were performed within a custom program written in Interactive Data Language (IDL 6.3). Details of the program were provided in two recent studies (Chen et al., 2023; Morales et al., 2023). The program is also briefly described below.

In general, ROI locations within a cell were chosen based on the appearance of obvious increases in fluorescence of the Ca^2+^ indicator. The mean intensity of each ROI was reported vs time as a Fluorescence at time *t* minus Initial Fluorescence / Initial Fluorescence, (F*_t_*-F_0_)/F_0_ or ΔF/F. This value is multiplied by 100 to arrive at %ΔF/F. The amplitude, time duration, and time course details of the Ca^2+^ signal response to agonists was strongly dependent on agonist. We chose several features of the response to characterize the similarities and differences. Some of these features were affected by photon shot noise which adds a jittery white noise to the signal unrelated to the biological processes. To circumvent shot noise for those features, we generated a smoothed version of the response by convolution with a Gaussian “kernel”. The 1/e width of that kernel is chosen to be 100 time bins wide, meaning that the highest frequency components of the shot noise was essentially smoothed over that many bins. The smoothing procedure was applied three successive times to enhance the smoothing.

(a) *Maximum amplitude, total duration, and total spike area.* The IDL program detected the heights and time-bin locations of all of the maxima and minima on the smoothed data. The highest amplitude achieved during the smoothed response was reported, measured above the lowest minimum, defined as a “baseline”. The total duration is measured was the number of time bins between the very first and the very last of the detected minima. The total area underneath the (unsmoothed) raw fluorescence vs time curve was computed as the total of the heights in each time bin above the baseline in that total duration window.

(b) *Spikes.* The response often consisted of a series of distinct sharp maxima (“spikes”), due to separate Ca^2+^ flashes possibly originating at different times at either the same or different locations within any single ROI. For each maximum, we establish a local baseline because that maximum often appeared above a broader base intensity variation or partially overlaps with other maxima. The local baseline was defined as the straight line that connects the two minimum (just before and just after) each maximum. If the height of a maximum above its local baseline (at the time bin of the maximum) was a certain specified fraction of the height of its local baseline, we declared that maximum to be a “spike”. In this way, only the most distinctly prominent of the maxima survive this “spike” test. That specified fraction here was varied between 0.2-0.25. The time duration of each spike was recorded as the number of time bins between its two surrounding minima. The “area” of a spike was the sum of the heights (with baseline subtracted) of each time bin between the two surrounding minima. The program reports: 1) total number of maxima and its subset number of spikes; 2) the height of each spike and its average over all the spikes; and 3) the area of each spike, the average area over all the spikes, and the total area of the spikes.

#### Imaging and image analysis of calcium puffs

TIRF images were captured at 80 frames per second with a 16-bit depth, utilizing 2 × 2 binning for a 128 × 128 binned pixel specimen field (one binned pixel equivalent to 320 nm). Ca^2+^ puff events were visually identified and ROIs selected using the Time Series Analyzer v3.0 plugin in the Fiji software. Calcium puff analysis was performed on OriginPro 2023b (Origin Lab Corporation) using the “Peak Analyzer” function. After a baseline was established, peaks were automatically identified, then filtered by height (threshold 15%) and fit to a Gaussian function.

### Electrophysiology

Voltage clamp recordings were made in the perforated patch clamp configuration using a Multiclamp 700b patch clamp amplifier (Molecular Devices). Data acquisition was performed using an Axon DigiData 1550B digitizer with the Clampex 11.1 software (Molecular Devices) and sampled at 2 kHz. All experiments were performed at 35°C on primary cell cultures maintained for 24-48 hrs using borosilicate glass pipettes constructed from 1.5 mm outer diameter (o.d) capillary glass tubing containing a filament (Warner Instruments, Hamden, CT). Pipettes were coated with Sylgard Elastomer (DOW Silicones Corp, Midland, MI) and fire polished to resistances of 1.5-3 MΩ. The intracellular recording solution consisted of (in mM): KCl (135), NaCl (8), MgCl_2_ (2), HEPES (20) with a pH of 7.2 with KOH and included 70-100 μg/mL of amphotericin B (Millipore-Sigma) added just prior to recording. External solution consisted of (in mM): NaCl (125), KCl (5.3), HEPES (10), MgCl_2_ (0.8), CaCl_2_ (10), Glucose (15) and a pH of 7.4 using NaOH. Cells were held at a membrane potential of -55 mV and evoked currents were measured during the bath application of either acetylcholine (100 nM) or PACAP (500 nM). In some experiments, the L-type Ca^2+^ channel blocker nifedipine (10 µM) was applied during PACAP application. Traces were analyzed using Clampfit 11.1 and peak current amplitudes were determined. To compensate for potential effects of cell size on current recordings, the peak amplitudes werr normalized to the corresponding whole cell capacitance measured during the experiment (pA/pF). All statistical analysis was performed using GraphPad. Data was expressed as mean ± SEM. Comparisons between two groups were analyzed by a student’s paired t-test. Statistical significance was considered where p value < 0.05.

### Statistical Analysis

GraphPad Prism was used for all statistical analysis. Curve fitting was performed on OriginPro 2023b or IDL. The distribution of values for a particular data set were first assessed for normality with a Shapiro-Wilk test. Differences in means of normally distributed data were subsequently compared using a Student’s t-test with Welch’s correction. A Mann-Whitney test was used to compare data sets whose individual values were not normally distributed (Dwivedi et al., 2017). For multiple comparisons, either a One-way ANOVA (normally distributed data) with Tukey’s test or Kruskal-Wallis with Dunn’s test was used to compare data sets. Scatter plots report means ± standard error of the mean (SEM) for normally distributed data, or medians with 25^th^ and 75^th^ percentile values for non-normally distributed data. Exact p values are reported except when differences between groups are not significant (i.e., p > 0.05). Within figures, the symbol ns means p > 0.05, * means p ≤ 0.05, ** means p ≤ 0.01, *** means p ≤ 0.001, and **** means p ≤ 0.0001. P values are reported to 1 significant digit. In a few cases, because of rounding, the reported p value does not match the asterisk scheme above. All other numerical values are reported to 2 or 3 significant digits.

## Results

### PACAP-stimulated Ca^2+^ signals require extracellular Ca^2+^

The major goal of this study was to determine the mechanisms by which PACAP causes changes in cytosolic Ca^2+^ that are important for secretion. As a first step towards this goal, the effects of PACAP stimulation, in the presence and absence of external Ca^2+^, were characterized. For these studies, chromaffin cells were transfected to express a GCaMP5G indicator that was tethered to the plasma membrane by Lck (Shigetomi et al., 2010). In this way, the Ca^2+^ signal in the region of the cell best imaged by TIRF is greatly accentuated. In the example shown in **Figure 1A**, a chromaffin cell expressing Lck-GCaMP5G was stimulated with 500 nM PACAP in a physiological solution containing 2.2 mM Ca^2+^. This caused a fluctuating fluorescent signal, increasing and decreasing with a variable amplitude and time course (**Figure 1B**). While the specific characteristics of the PACAP-stimulated Ca^2+^ signals were heterogeneous, increasing concentrations of PACAP generally produced spikes with greater amplitudes (**Figure S1A**).

**Figure 1.**
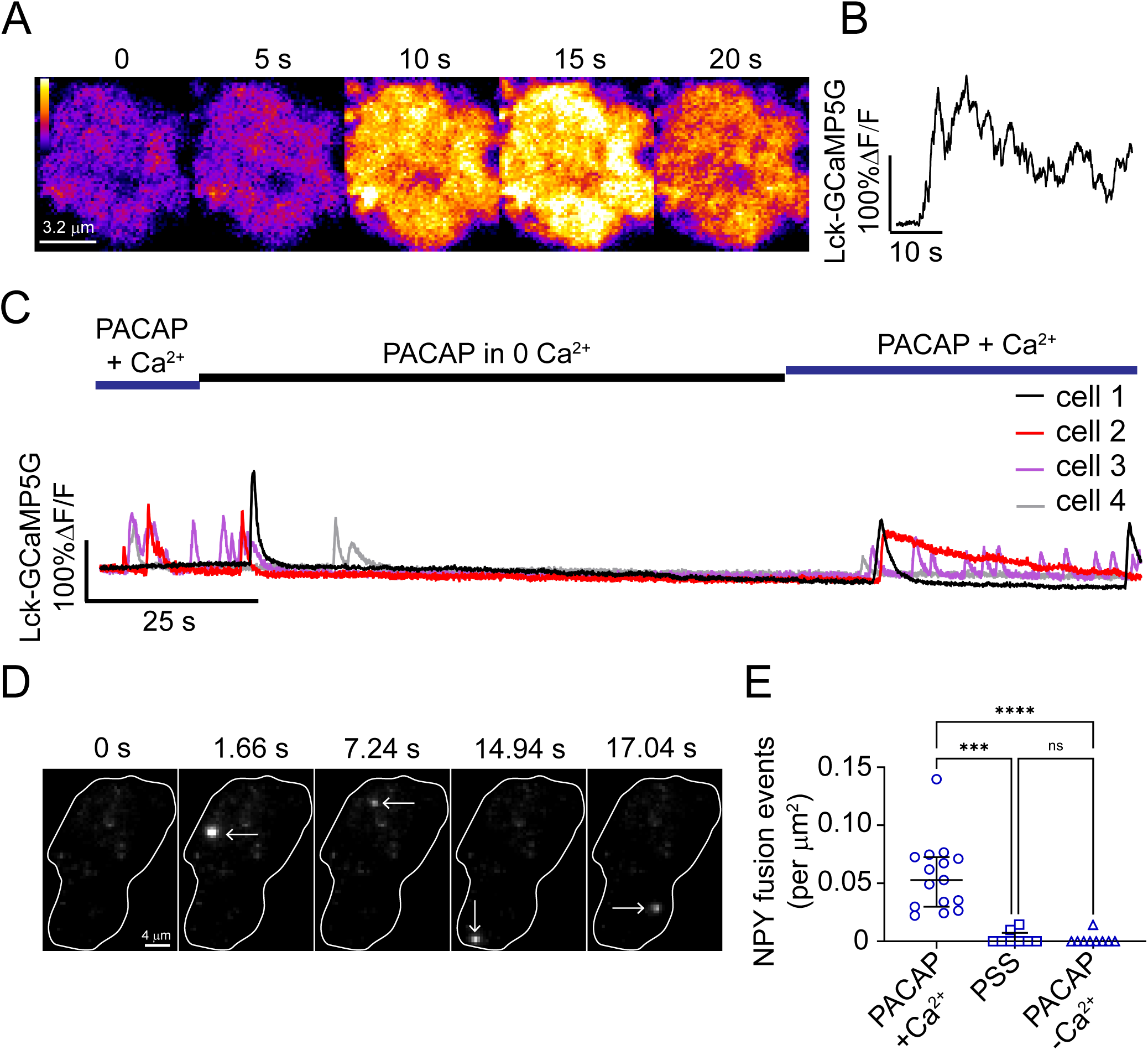
PACAP-stimulated cytosolic Ca^2+^ elevations and secretion required extracellular Ca^2+^. **A**. Representative pseudocolour images of a cell expressing Lck-GCaMP5G prior to stimulation (time 0) and during stimulation with 500 nM PACAP. **B**. The percentage change in fluorescence (%ΔF/F) versus time record for the cell in **A**. **C**. %ΔF/F versus time records show external Ca^2+^ was required for PACAP to stimulate cytosolic Ca^2+^ transients. 4 representative traces are shown (n = 7 total). **D.** Cell expressing NPY-pHluorin was stimulated by 500 nM PACAP. The outline of the cell footprint is indicated in white. Arrows show the location of individual NPY fusion events in the time series. **E**. PACAP-stimulated exocytosis required extracellular Ca^2+^. Cells were either stimulated by PACAP with (+) extracellular Ca^2+^ (n=15), PSS (n=9), or PACAP without (-) Ca^2+^ (n=9). A Kruskal-Wallis test was performed for significance testing. PSS vs PACAP+Ca^2+^, p = 0.0003; PACAP-Ca^2+^ vs PACAP+Ca^2+^, p = 3×10^-5^. Scatter plots show medians ± interquartile range.

We next tested the effect of removing external Ca^2+^ in the solution bathing the cell during the PACAP stimulation period. In these experiments, 4 examples of which are shown, cells were initially stimulated with PACAP in normal Ca^2+^, then switched to a stimulation medium in which Ca^2+^ was absent (**Figure 1C**). The fluorescent cytosolic Ca^2+^ signals, initially robust and possessing the characteristics associated with PACAP stimulation (Chen et al., 2023; Morales et al., 2023), disappeared once external Ca^2+^ was removed. When normal [Ca^2+^]_ex_ was returned to the stimulation solution, the fluorescent signals promptly returned. The reverse experiment was also performed, where Ca^2+^ was initially absent from the stimulation medium (**Figure S1B**). Here too, it was only in normal [Ca^2+^]_ex_ that PACAP caused increases in Lck-GCaMP5G fluorescence (**Figure S1B**).

Without external Ca^2+^, Lck-GCaMP5G responses to PACAP stimulation were not detected in chromaffin cells. Thus, it also seemed unlikely that under these conditions PACAP would cause secretion. To test this idea, the secretory response of cells stimulated by PACAP in the presence and absence of external Ca^2+^ was compared. Secretion was visualized by monitoring the fluorescence of overexpressed neuropeptide Y (NPY) pHluorin (Anantharam et al., 2010; Miesenbock et al., 1998). Sudden increases in NPY-pHluorin fluorescence on the footprint indicated the time and location of a fusion event (**Figure 1D**; see arrows). While fusion events were readily observed in cells stimulated by a PACAP medium containing external Ca^2+^, few events, if any were observed when external Ca^2+^ was absent. In fact, the probability of fusion in a Ca^2+^-free PACAP medium was indistinguishable from the likelihood of spontaneous fusion (i.e., where cells were exposed only to physiological saline, PSS; **Figure 1E**).

### ER Ca^2+^ depletion inhibits PACAP-evoked Ca^2+^ signals and exocytosis

The process of PACAP-stimulated exocytosis depends critically on external Ca^2+^ (**Figure 1**). Thus, it has been widely assumed that PACAP-stimulated secretion is mediated by Ca^2+^ entry pathways, especially voltage-gated channels, located on the plasma membrane of chromaffin cells and PC12 cells (Hill et al., 2011; Kuri et al., 2009; O’Farrell and Marley, 1997; Osipenko et al., 2000; Tanaka et al., 1996). However, if PACAP relied exclusively on voltage-gated channels, the secretory response would eventually run down as channels inactivated and Ca^2+^ influx was restricted; this does not seem to be the case (Chen et al., 2023; Morales et al., 2023). These observations suggested to us that PACAP must harness additional Ca^2+^ sources to cause secretion. We pursued this idea by focusing first on the possible involvement of internal Ca^2+^ stores (Mustafa et al., 2007; Tanaka et al., 1996).

The properties of a PACAP Ca^2+^ signal that is primarily dependent on external Ca^2+^ should not be altered by Ca^2+^ store depletion. One way in which internal Ca^2+^ stores can be rapidly depleted is by exposing cells to caffeine. Once inside the cell, caffeine is known to activate ryanodine receptors (RyRs), possibly by increasing receptor open probability, and cause Ca^2+^ release from the ER into the cytosol (Eisner et al., 2017; Kong et al., 2008; West and Williams, 2007). Indeed, a long-lasting increase in cytosolic Ca^2+^ was observed in chromaffin cells exposed to caffeine for 3 min (**Figures 2A-2B**; **Figure S1C**). The perfusion of PACAP onto chromaffin cells immediately after caffeine failed to have much of an effect on cytosolic Ca^2+^. However, if cells were washed for 5 min in physiological saline and then stimulated by PACAP, a substantial increase in cytosolic Ca^2+^ was again observed (**Figures 2A**-**2B**). The mean “pre-wash” spike area in cells stimulated by PACAP was 480 ± 82; the “post-wash” spike area in cells stimulated by PACAP was 2500 ± 780 (p = 0.04) (**Figure 2C**).

**Figure 2.**
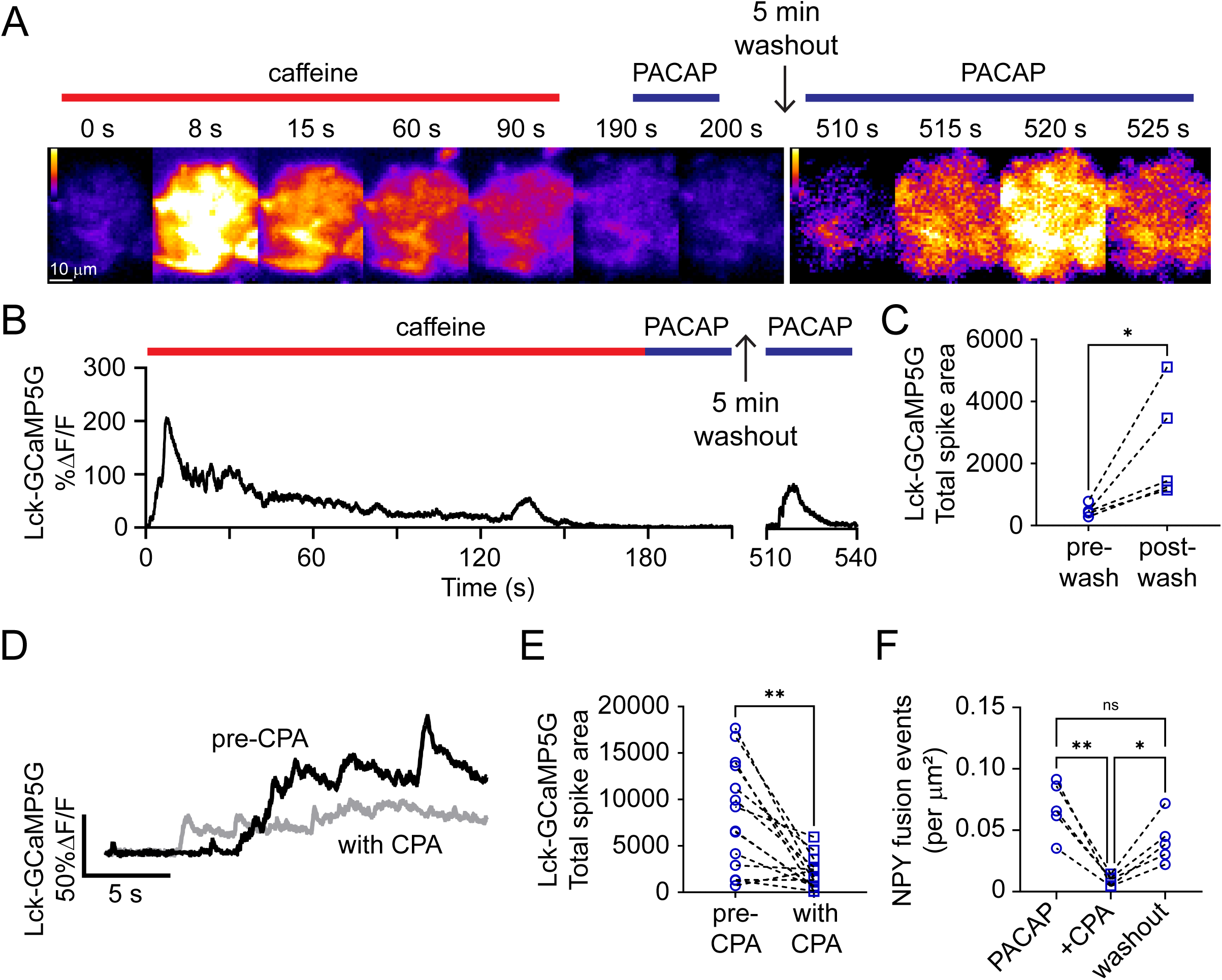
Depletion of ER Ca^2+^ stores reduced PACAP-stimulated Ca^2+^ signals and secretion in chromaffin cells. **A**. A cell expressing Lck-GCaMP5G was exposed to 40 mM caffeine for 180 s and then immediately stimulated by 500 nM PACAP for 30 s. After a 5 min wash with PSS, the cell was again stimulated by 500 nM PACAP. PACAP was ineffective at stimulating Ca^2+^ signals prior to the PSS wash. **B**. %ΔF/F versus time record for the cell in **A**. **C**. The total Ca^2+^ signal spike area was quantified for cells stimulated with PACAP before or after PSS wash. A paired Student’s t-test used to assess significance (p = 0.04). **D**. %ΔF/F versus time records show a representative Lck-GCaMP5G fluorescent response to PACAP before (black line) and after (grey line) 10 μM CPA treatment. **E**. The total Ca^2+^ signal spike area for n=14 cells stimulated with PACAP before or after 10 μM CPA treatment. Difference in means is statistically significant with paired t-test (p = 0.001). **F**. NPY fusion events per unit area were measured under three different conditions: 1) in response to 500 nM PACAP stimulation before CPA treatment; 2) after CPA (10 μM) treatment; and, 3) following the washout of CPA. Each cell (n=6) was first stimulated with PACAP and then incubated with CPA for 5 min. The cell was then stimulated again with PACAP in the continued presence of CPA. Finally, the CPA was washed out with PSS for 5 min, and the cell was stimulated with PACAP again. Friedman’s test with Dunn’s multiple comparisons were used to assess significance. PACAP vs PACAP+CPA, p = 0.004; PACAP+CPA vs PACAP washout, p = 0.04).

We next examined the effect of blocking Ca^2+^ loading into the ER. Application of the sarco/endoplasmic Ca^2+^ ATPase (SERCA) inhibitor, cyclopiazonic acid (CPA), significantly reduced the spike area of PACAP-stimulated Ca^2+^ signals from a mean of 8200 ± 1500 to 2000 ± 460 (p = 0.001) (**Figures 2D-2E**). CPA treatment also impacted exocytosis. The number of NPY-pHluorin fusion events per unit area fell from a mean of 0.073 ± 0.01 to 0.013 ± 0.004 (p = 0.005). After a 5 min wash with PSS, exocytosis returned to pre-CPA levels (**Figure 2F**).

### PACAP causes Ca^2+^ release from the ER

The depletion of ER Ca^2+^ disrupted both PACAP-evoked Ca^2+^ signals and exocytosis (**Figure 2**). This suggested to us that a major part of the process by which PACAP causes exocytosis involves Ca^2+^ mobilized from the ER. We therefore measured ER Ca^2+^ dynamics directly using an ER lumen-targeted GCaMP protein (ER-GCaMP6-150) (de Juan-Sanz et al., 2017). ER-GCaMP6-150 expression revealed the cellular ER to consist of an elaborate network of tubules extending from the perinuclear region to the plasma membrane of chromaffin cells (**Figure 3A**). Based on this pattern of expression, we predicted that, should PACAP cause changes in ER Ca^2+^ at or near the plasma membrane, they would be readily observable in TIRF (**Figure 3B**). Indeed, imaging experiments showed that PACAP caused small, but consistent, decreases in ER-GCaMP6-150 fluorescence (PACAP+Ca^2+^ mean -9.9 ± 1.8 %ΔF/F, n = 11) (**Figures 3C**-**D**). A representative %ΔF/F versus time record is shown in **Figure 3C**. **Figures 3C-3D** show that when PACAP is used to stimulate cells in the absence of Ca^2+^, the decrease in ER-GCaMP6-150 fluorescence was more pronounced. The most parsimonious explanation for this observation is that PACAP-stimulated ER Ca^2+^ release is usually balanced by a reloading of the ER that relies on external Ca^2+^. Thus, in conditions where external Ca^2+^ is not available (i.e., the PACAP minus Ca^2+^ condition), PACAP continues to cause ER Ca^2+^ release without appreciable ER Ca^2+^ reloading. This is reported visually as a substantial fall in ER-GCaMP6-150 fluorescence (mean -73 ± 3.0% ΔF/F; n = 11) that occurs within 30 s. Note, ER-GCaMP6-150 fluorescence did not appreciably change when Ca^2+^ was removed from the extracellular medium in unstimulated cells (**Figure 3E**).

**Figure 3.**
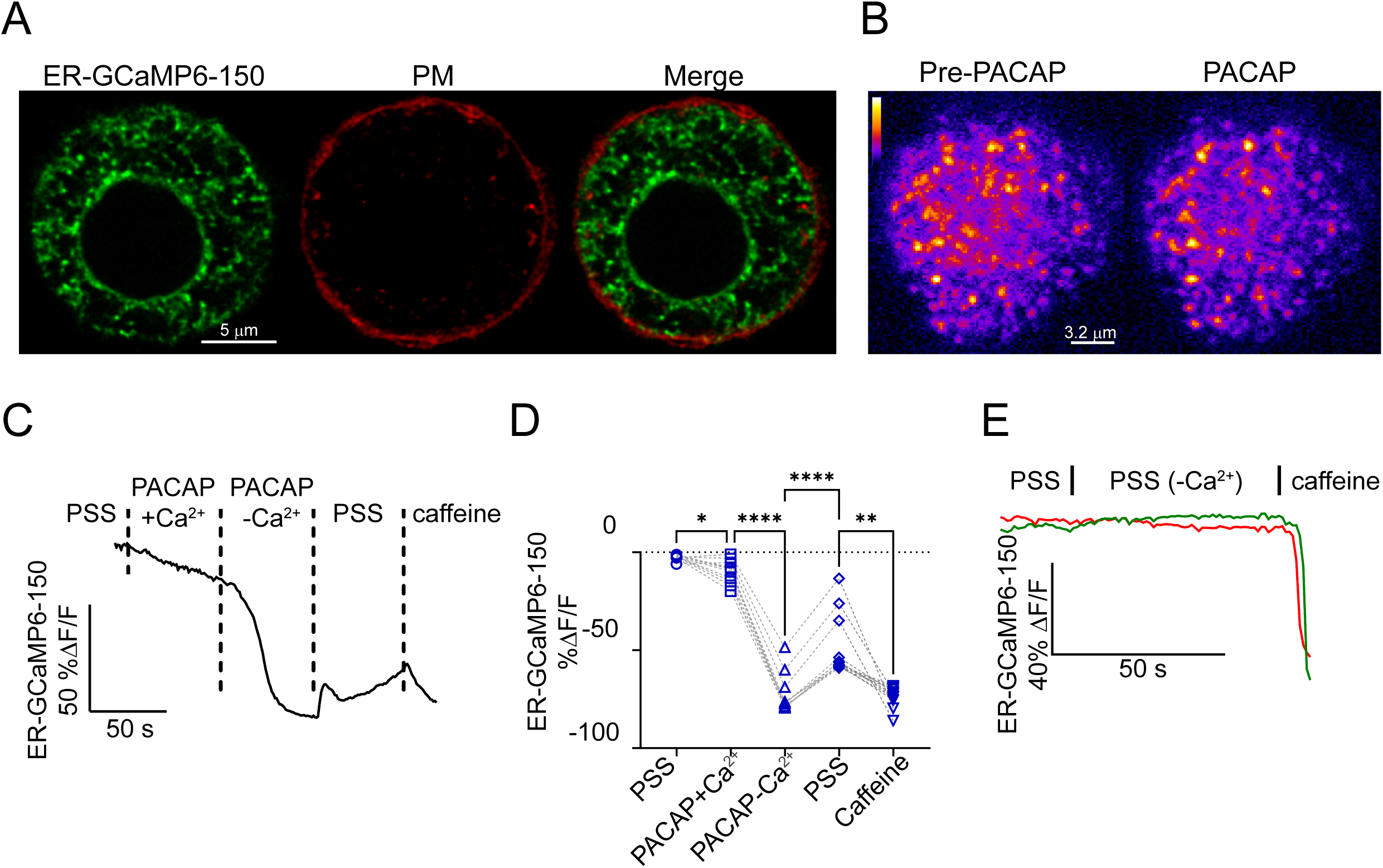
PACAP elicited ER Ca^2+^ release in chromaffin cells. **A**. Representative confocal images of a chromaffin cell expressing ER-GCaMP6-150. The plasma membrane (PM) was stained with CellMask™ Deep Red. **B**. Representative TIRF images of a cell expressing ER-GCaMP6-150 prior to stimulation and after stimulation with 500 nM PACAP. **C**. Exemplar record of an ER-GCaMP6-150 response to the stimulation paradigm indicated. When extracellular [Ca^2+^]_ex_ was reduced to ∼ 0 mM, the PACAP-stimulated decline in ER-GCaMP6-150 fluorescence was accelerated. **D**. Scatter plots showing individual maximum decreases in ER-GCaMP6-150 fluorescence in response to the conditions indicated (n=15). Data were compared for significance using a one-way ANOVA with Tukey’s multiple comparison’s test. PSS vs PACAP+Ca^2+^, p = 0.02; PACAP+Ca^2+^ vs PACAP-Ca^2+^, p = 3×10^-10^; PACAP-Ca^2+^ vs PSS, p = 1×10^-6^; PSS vs caffeine, p = 0.009. Not all comparisons are shown for clarity. **E**. PACAP stimulation was required for ER Ca^2+^ release. In cells exposed sequentially to PSS + extracellular Ca^2+^ and PSS minus (-) extracellular Ca^2+^, little to no change in ER-GCaMP5G fluorescence was observed (2 examples are shown; representative of 7 cells).

### PACAP-stimulated Ca^2+^ signals require Ca^2+^ influx through an L-type, nifedipine-sensitive <u>channel</u>

The experiments in **Figure 3** suggest that maintenance of ER Ca^2+^, even in the short term, is dependent on Ca^2+^ entry. We previously showed that PACAP caused a small membrane depolarization in chromaffin cells of between 2-10 mV and increased its spiking activity (Morales et al., 2023). One way in which this might happen is via activation of a low-voltage activated, L-type Ca^2+^ channel (Marcantoni et al., 2008; Marcantoni et al., 2010). To test this idea, cells expressing Lck-GCaMP5G were stimulated with PACAP, and then exposed sequentially to 1 and 10 μM nifedipine, which blocks L-type channels. The PACAP-stimulated Ca^2+^ signals were reduced by nifedipine in a dose-dependent manner (**Figures 4A-4D**). Nifedipine, applied at a concentration of 10 μM, strongly inhibited the fusion of NPY-pHluorin-labeled granules (**Figure 4E**).

**Figure 4.**
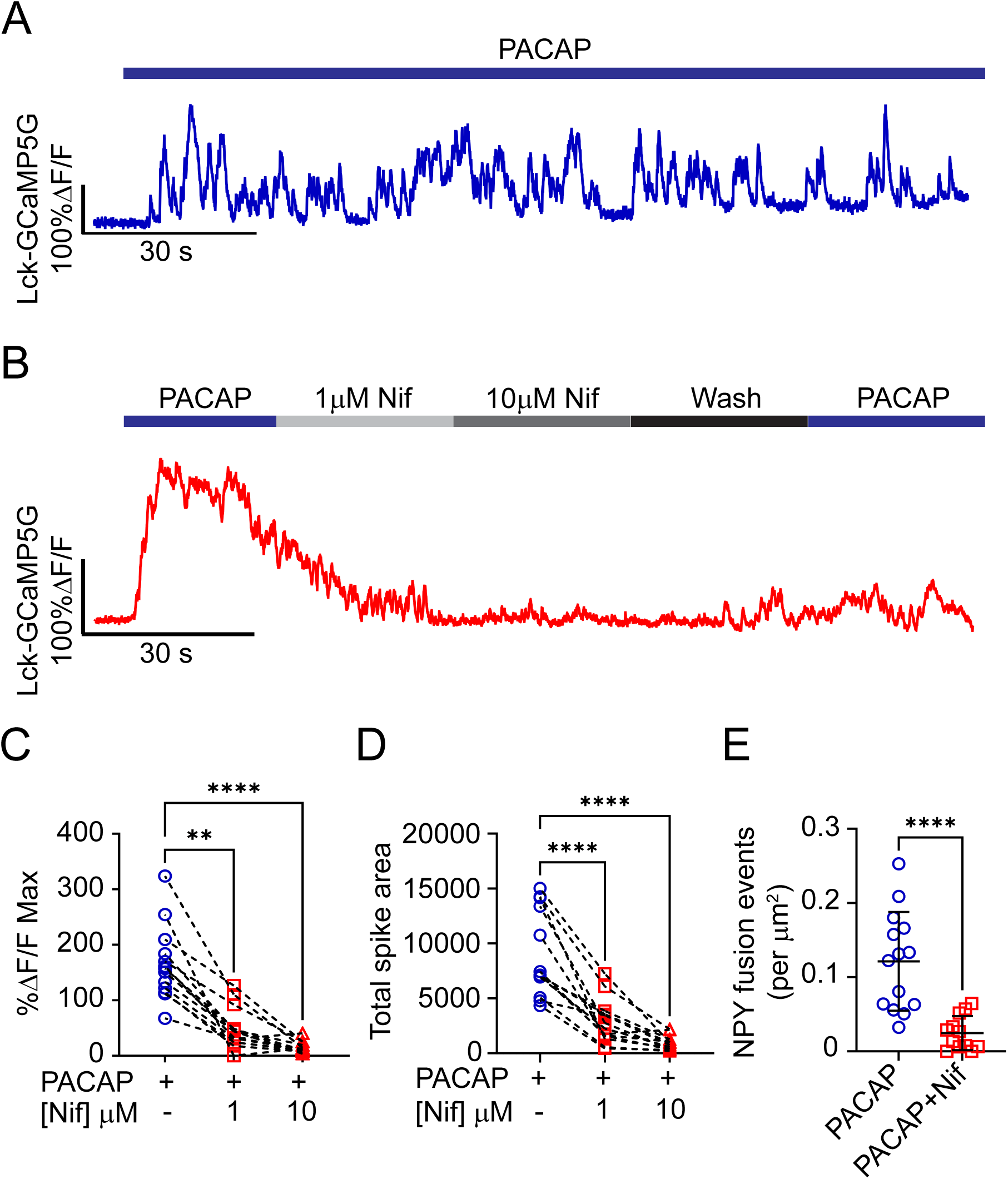
The L-type Ca^2+^ channel blocker, nifedipine, reduced PACAP-evoked Ca^2+^ transients and secretion in chromaffin cells. **A**. Representative %ΔF/F versus time trace for a chromaffin cell expressing Lck-GCaMP5G and stimulated with 500 nM PACAP for 150s. **B**. Representative %ΔF/F versus time trace for a sample chromaffin cell showing Ca^2+^ responses to 500 nM PACAP before and after application of 1 or 10 μM nifedipine. Washout of nifedipine partially restored PACAP response. **C**. Maximum %ΔF/F for chromaffin cells (n=13) stimulated with 500 nM PACAP alone or with two different nifedipine concentrations, Kruskal-Wallis test with Dunn’s multiple comparisons were used to assess significance. PACAP vs. PACAP+1 μM nifedipine, p = 0.004; PACAP vs. PACAP+10 μM nifedipine, p = 7×10^-6^. **D**. PACAP-stimulated Ca^2+^ spike area was reduced by nifedipine in a dose-dependent manner. Kruskal-Wallis with Dunn’s multiple comparisons test used for significance testing. PACAP vs. PACAP+ 1 μM nifedipine, p = 8×10^-9^; PACAP vs. PACAP+10 μM nifedipine, p = 3×10^-12^. **E**. NPY secretion in the absence (n=14) or presence (n =12) of nifedipine. Differences in the means are statistically significant (Student’s t-test), p = 7×10^-5^.

To examine whether PACAP application causes the appearance of nifedipine-sensitive current, perforated patch-clamp electrophysiology was performed (**Figure S2A**). The results show PACAP did activate an inward current was much smaller and more rapidly desensitizing than the current measured after ACh application (mean 2.7 pA/pF vs 68 pA/pF). Importantly, the PACAP-stimulated current was significantly reduced by nifedipine application (mean 3.9 pA/pF to 0.9 pA/pF) (**Figures S2A-S2D**). In contrast to nifedipine, N- (ω-conotoxin) and P/Q-type channel (ω-agatoxin) blockers had no appreciable effect on either the amplitude or the area under the curve of the Ca^2+^ signals evoked by PACAP (**Figures S3A-S3D**).

### ER Ca^2+^ release likely depends on IP3 receptor activation

Our results show that PACAP mobilizes Ca^2+^ as part of the mechanism for exocytosis (**Figure 2**). We speculated that the pathway by which this occurs is likely to involve either RyRs or IP3Rs. Both types of receptors gate the movement of Ca^2+^, but only RyRs have been suggested to be involved in the pathway of PACAP-stimulated secretion in chromaffin cells (Mustafa et al., 2010; Payet et al., 2003; Tanaka et al., 1996). We systematically assessed the involvement of RyRs and IP3Rs by employing pharmacological agents that antagonized their function. Because the goal was to monitor how receptor inhibition impacts near-membrane Ca^2+^ levels at or near sites of exocytosis, Lck-GCaMP5G was used as the Ca^2+^ indicator. These experiments revealed that, while ryanodine had little effect on the properties of PACAP-stimulated Ca^2+^ signals, 5 μM Xestospongin C (Payet et al., 2003) – an antagonist of IP3Rs – inhibited total spike area and amplitude (**Figures 5A-5B**). Next, we investigated the effect of inhibiting IP3R function on exocytosis. Cells expressing NPY-pHluorin were stimulated with PACAP after a brief incubation with Xestospongin C. **Figure 5C** shows that PACAP-stimulated fusion events were significantly inhibited by Xestospongin C (**Figure 5C**).

**Figure 5.**
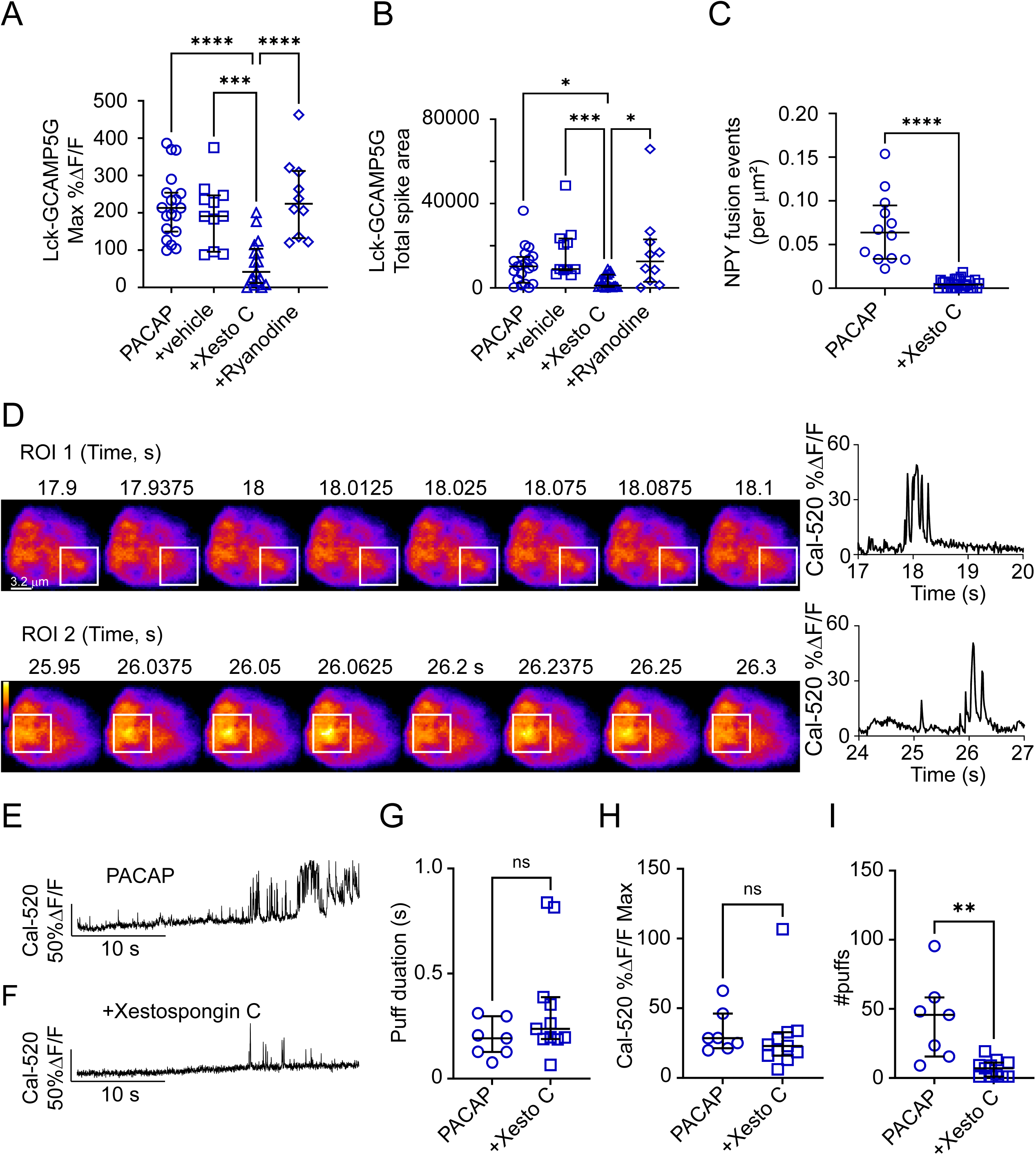
IP3Rs were activated by PACAP. **A**. An IP3R inhibitor, Xestospongin C (Xesto C) reduced the amplitude of PACAP-evoked Ca^2+^ transients. Cells (for **A** and **B**) were perfused with either PACAP (500 nM) alone, PACAP+DMSO (0.1%), PACAP+Xesto C (5 μM), or PACAP+ryanodine (100 μM). DMSO, Xesto C, and ryanodine were also added to the bath at least 1 min prior to perfusion at the indicated concentration. n=19 for PACAP, n=11 for PACAP+vehicle, n=17 for PACAP+Xesto C, n =10 for PACAP+ryanodine. One-way ANOVA was used for significance testing. PACAP vs. PACAP+Xesto C, p = 7×10^-6^; PACAP+vehicle vs. PACAP+Xesto C, p = 0.0007; PACAP+ryanodine vs. PACAP+Xesto C, p = 2×10^-5^. **B**. Xestospongin C reduced total area of Ca^2+^ spikes. PACAP vs. PACAP+Xesto C, p = 0.02; PACAP+vehicle vs. PACAP+Xesto C, p = 0.0009; PACAP+ryanodine vs. PACAP+Xesto C, p = 0.01. **C**. The secretory response to PACAP (500 nM) was disrupted by Xesto C (5 μM), n =12. Differences in the means are statistically significant (unpaired Student’s t-test), p = 6×10^-6^. **D**. Two examples of Ca^2+^ puffs (boxed) imaged with Cal-520 are shown with corresponding intensity versus time records. **E** and **F**. Examples of Ca^2+^ puffs in cells stimulated by PACAP or PACAP + Xestospongin C. **C**. Xestospongin C was applied as in **A** and **B**, above. **G and H.** Data from multiple ROIs in each cell were averaged and are presented as a cell average (PACAP, n=7; +Xestospongin C, n = 11). No statistically significant differences in puff duration or amplitude were observed in chromaffin cells stimulated with PACAP or PACAP+Xestospongin C (Mann-Whitney). Scatter plots show median ± interquartile range. **I**. The number of puffs appearing in cells exposed to PACAP+Xestospongin C was substantially lower than in cells exposed to PACAP alone (Student’s t-test; p = 0.001). Scatter plots show mean ± SEM.

To determine whether the inhibition caused by Xestospongin C was specific to the PACAP stimulation pathway, we also exposed chromaffin cells expressing Lck-GCaMP5G to DMPP – a nicotinic receptor agonist that also causes secretion (Pothos et al., 2002). Interestingly, Xestospongin C, at the 5 μM concentration that strongly reduced PACAP-stimulated Ca^2+^ signals, had no effect on Ca^2+^ signals stimulated by DMPP or on DMPP-triggered exocytosis (**Figures S4A-C**).

IP3R-mediated Ca^2+^ signals are organized at multiple levels – from “blips” to “waves” – thereby encoding different types of information based on how strongly the cell is stimulated (Foskett et al., 2007). The coordinated opening of a few IP3R channels causes Ca^2+^ to be released in “puffs” (Foskett et al., 2007; Lock and Parker, 2020). Ca^2+^ puffs are thus considered to be the “building blocks” of much larger global Ca^2+^ signals (Lock et al., 2018; Lock and Parker, 2020; Marchant and Parker, 2001; Parker et al., 1996). To image IP3R-mediated Ca^2+^ puffs, chromaffin cells were loaded with Cal520-AM (Arige et al., 2021). Cells normally maintained in 2.2 mM Ca^2+^ were briefly switched to a solution containing ∼ 0 mM Ca^2+^ and 300 μM EGTA for 30 s. During this time window, cells were directly stimulated with PACAP to elicit IP3 production and generate Ca^2+^ puffs. The 0 mM Ca^2+^ solution was necessary to prevent PACAP from causing global Ca^2+^ waves that might obscure puff detection.

Examples of PACAP-stimulated Cal520 Ca^2+^ signals are shown in **Figure 5D**. Regions of interest in which Ca^2+^ puffs emerge are indicated by white boxes. Intensity versus time records are shown to the right of the image panels. Ca^2+^ puffs were also imaged in the presence of Xestospongin C (compare **Figure 5E** to **Figure 5F**). The major effect of Xestospongin C was not to reduce either the duration or amplitude of the Ca^2+^ puffs (**Figures 5G-5H**), but rather to reduce the frequency with which they occur (**Figure 5I**). Therefore, it appeared that Xestospongin C inhibited PACAP-stimulated Ca^2+^ signals in chromaffin cells by disrupting the activity of IP3Rs.

We previously showed that PACAP-stimulated secretion requires PLCε (Morales et al., 2023). Thus, in cells lacking PLCε expression, PACAP should not cause IP3 production or stimulate Ca^2+^ puffs. To test this idea, we again imaged Ca^2+^ puffs in chromaffin cells, this time in PLCε KO cells that were exposed to PACAP. KO cells stimulated by PACAP exhibited very few puffs (**Figures 6A-6B**). However, when PLCε was overexpressed in KO cells (i.e,. the “rescue” condition), many more puffs were observed. This is shown both in the exemplar trace (**Figure 6A**) and in the graph (**Figure 6B**). These experiments show that PACAP-stimulated Ca^2+^ puffs require PLCε. This is presumably because IP3 production and IP3R function depend on the stimulation of PLCε activity.

**Figure 6.**
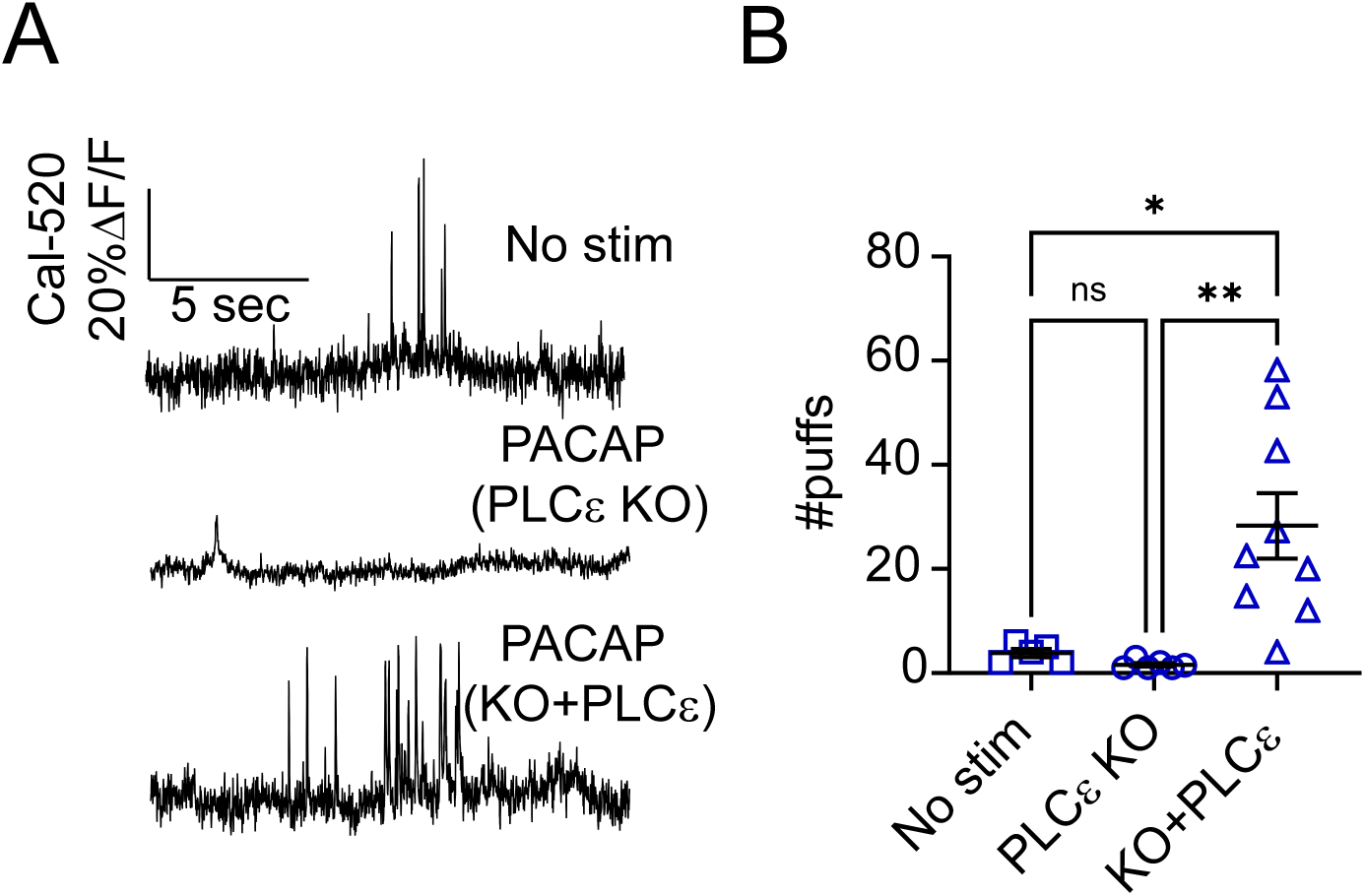
PACAP-stimulated Ca^2+^ puffs require PLCε expression. **A**. Examples of Ca^2+^ puffs in unstimulated cells (top; n=5) and PACAP-stimulated PLCε KO cells (middle; n=6) or KO cells in which PLCε was overexpressed (bottom; n=9). **B**. Scatter plots show means ± SEM. Differences in the means were statistically significant (Brown-Forsythe and Welch ANOVA test). No stim vs. KO+PLCε, p = 0.01; PLCε KO vs. KO+PLCε, p = 0.008).

### Chromaffin cells express multiple IP3R isoforms with type 1 being most abundant

Three isoforms of IP3Rs have been discovered, each encoded by a separate gene (Foskett et al., 2007). We first performed qPCR to assess which of these is expressed in mouse chromaffin cells. We identified transcripts for all three IP3Rs, with the type 1 isoform being the most highly expressed (**Figure 7A**). We also performed qPCR to probe RyR isoform expression and identified transcripts for receptor types 2 and 3 (**Figure 7A**). Western blotting was performed to confirm expression of the IP3R1 protein in chromaffin cells (**Figure 7C**). To rule out the possibility of reduced IP3R expression, or defective signaling upstream of PLCε (e.g., reduced cAMP production), qPCR and imaging studies were also performed on PLCε KOs.

**Figure 7.**
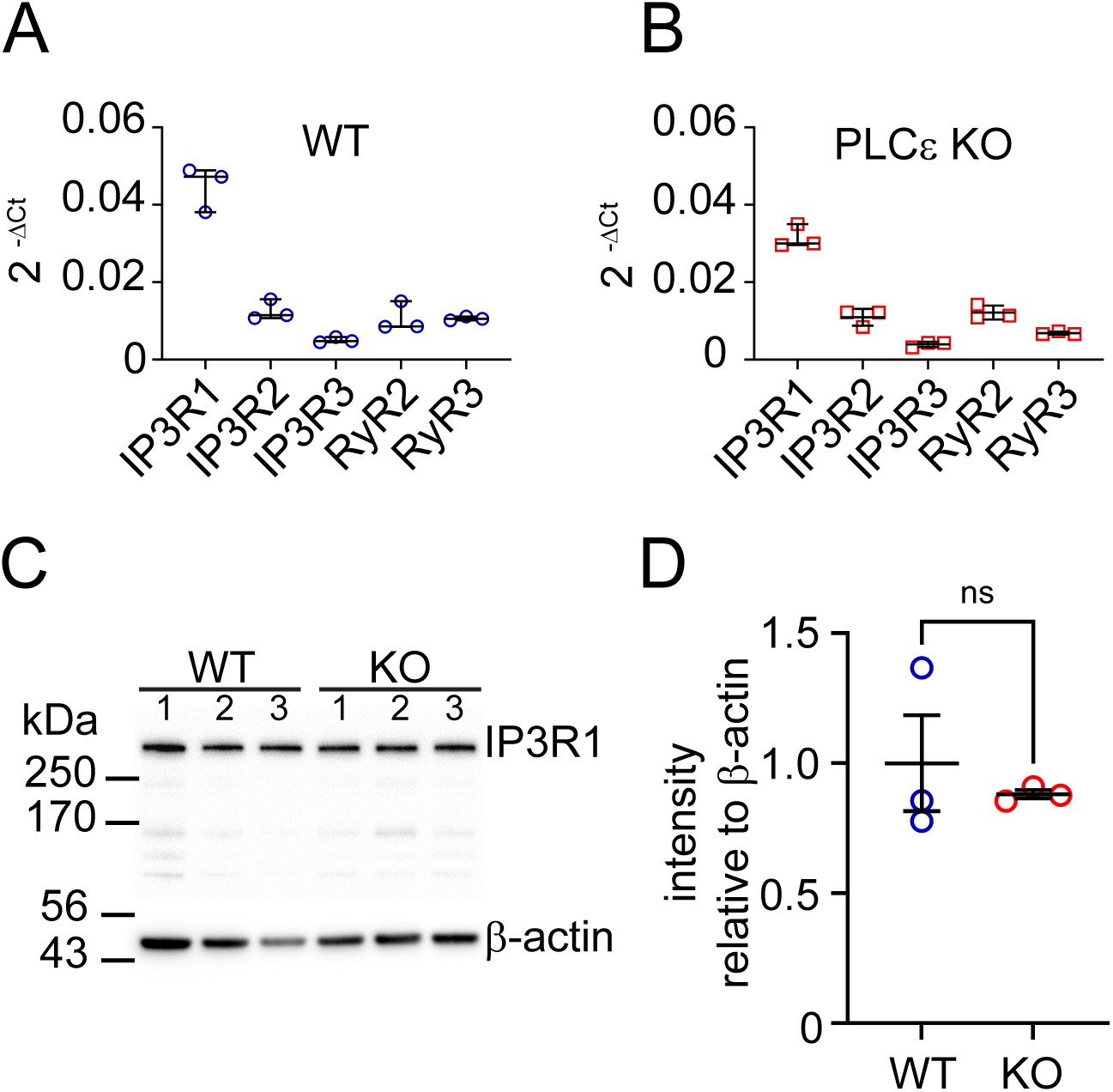
Chromaffin cells express multiple IP3R and RyR isoforms. **A**. Quantitative PCR showed chromaffin cells express all 3 of the IP3 receptor isoforms, but only 2 RyR isoforms. Mean transcript levels of IP3R1 (0.05 ± 0.003) were approximately 4-fold higher than IP3R2 (0.01 ± 0.002) and almost 10-fold higher than IP3R3 (0.005 ± 0.004). Expression levels were compared to β-actin. **B**. IP3R and RyR isoforms showed a similar pattern of expression in WT and PLCε KO cells. **C** and **D**. Western blot analysis comparing the protein expression levels of IP3R1 in WT and PLCε KO cells. The intensity of IP3R1 bands (WT vs KO) were quantified relative to β-actin and were not significantly different (Student’s t-test; p = 0.6).

First, we determined that the pattern of IP3R and RyR transcript expression was similar in WT and KO cells (**Figures 7A-7B**). Moreover, IP3R1 protein expression was appreciably different in WT and KO cells. The real-time production of cAMP and DAG, occurring upstream and downstream of PLCε, respectively, was also monitored optically in WT and PLCε KO cells (**Figures 5A-S5D**). The results show cAMP production to be normal in the KO. However, the formation of DAG was significantly inhibited in cells lacking PLCε expression (**Figures S5E-S5F**).

To probe the subcellular localization of the IP3R1 protein in chromaffin cells, confocal imaging was performed. Because we expected IP3R1 to be expressed on the ER, immunolocalization of the receptor was compared to that of the KDEL ER retention motif (Lewis and Pelham, 1992; Munro and Pelham, 1987). IP3R1 is widely distributed in the chromaffin cell, reflecting the broad expanse of the tubular ER network (evidenced by KDEL fluorescence) (**Figure 8A**). The subcellular expression profile of IP3R1 was evaluated two different ways.

**Figure 8.**
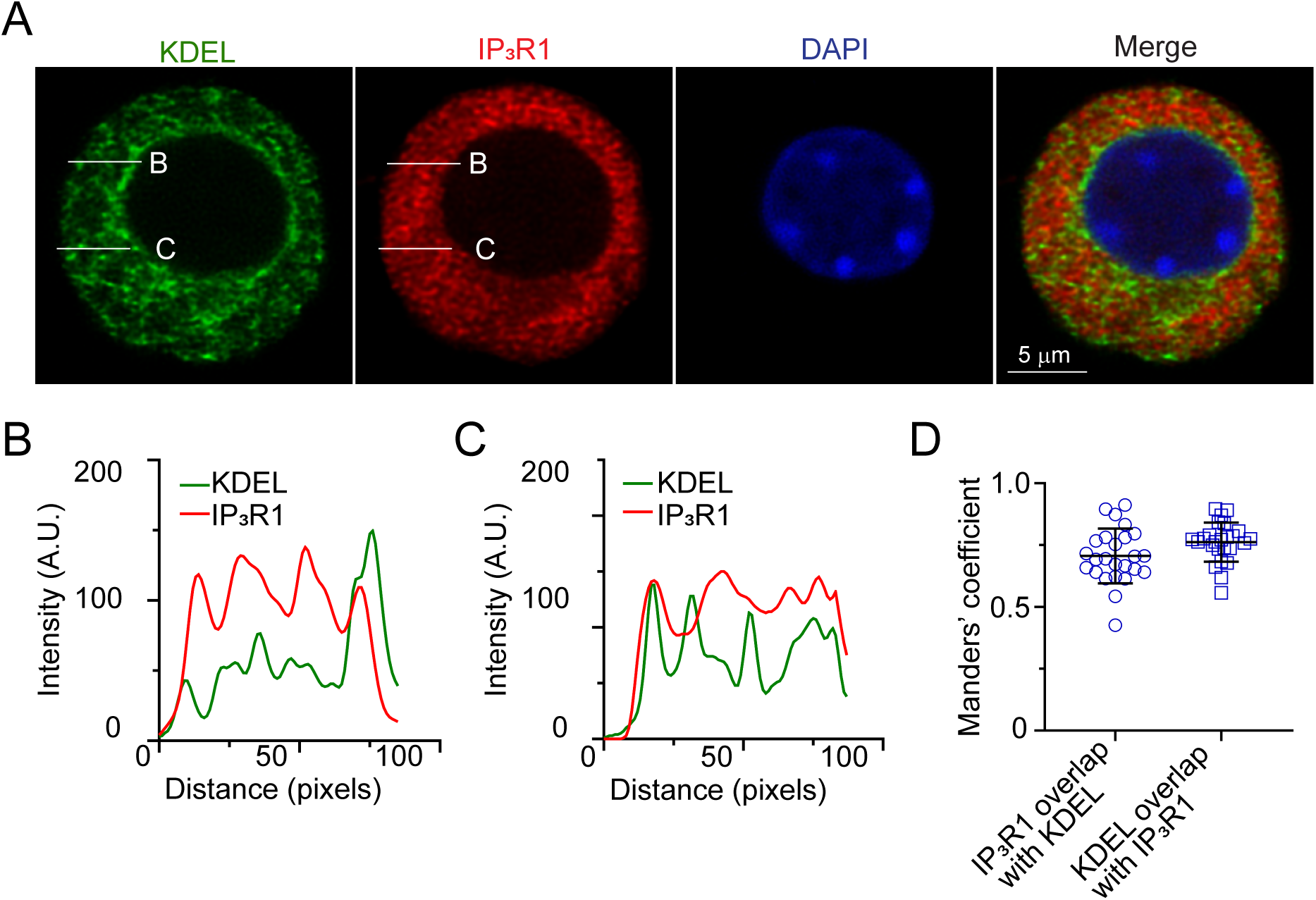
Colocalization of ER marker KDEL and IP3R1 in chromaffin cell. **A**. Confocal images showing immunostaining for the ER marker KDEL (green) and IP3R1 (red) in a chromaffin cell. Nuclei labeled with DAPI (blue). **B** and **C**. KDEL and IP3R1 intensities along the lines were similar. **D**. Manders’ coefficient analysis for the colocalization of KDEL and IP3R1 in n = 25 cells. Coefficient for IP3R1 overlap with KDEL = 0.7 ± 0.02; coefficient for KDEL overlap with IP3R1 = 0.8 ± 0.02.

First, lines were drawn along which the intensities of IP3R1 and KDEL were compared. **Figures 8B** and **8C** indicate a similar pattern of fluorescence along lines “B” and “C”. Second, Manders’ analysis was performed to measure the colocalization frequency of IP3R1 and KDEL fluorescence in chromaffin cells. The corresponding Manders’ coefficients averaged greater than 0.7, which is indicative of a high degree of overlap in the distribution of KDEL and IP3R1 proteins (**Figure 8D**) (Dunn et al., 2011).

### IP3R1 expression is required for normal responses to PACAP stimulation

PACAP failed to increase cytosolic Ca^2+^ or cause secretion in the PLCε KO. A similar effect was observed when IP3R activity was pharmacologically disrupted with Xestospongin C. Thus, the PACAP secretory response appears to depend critically on PLCε and downstream IP3R activity. To directly test this idea, we expressed an shRNA in chromaffin cells that targeted the IP3R1 transcript. **Figure 9A** shows IPR3R1 expression to be reduced by at least 60% in cells that took up the shRNA (**Figure 9B**). In fact, this is likely underestimating how much IP3R1 expression was depleted in cells. Because of the limited time cells were viable in culture, they were harvested for western blotting after only 2 days in the puromycin selection medium.

**Figure 9.**
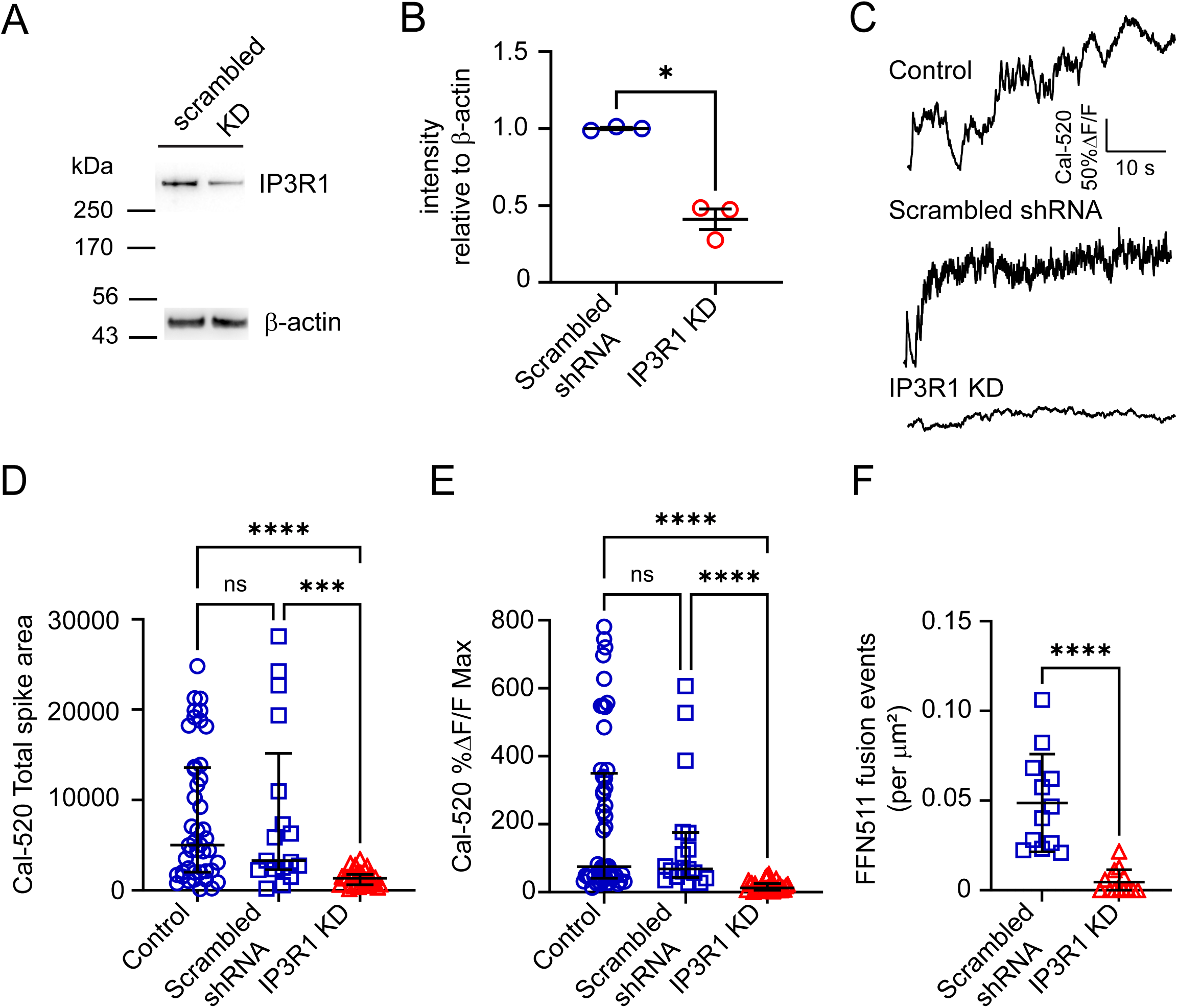
Knockdown of IP3R1 reduces PACAP-evoked Ca^2+^ signals and secretion. **A**. Western blot showing IP3R1 immunoreactive bands in control (scrambled shRNA) and IP3R1 KD samples. **B**. Intensity IP3R1 was normalized to β-actin expression. Relative intensities of bands (means ± SEM) of scrambled = 1 ± 0.007 and KD = 0.4 ± 0.06. Differences were statistically significant (Student’s t-test; p = 0.01). **C**. Representative traces of PACAP-stimulated Cal-520 fluorescence changes in control (untransfected) cells, scrambled shRNA, IP3R1 KD cells. **D**. Scatter plot of total spike area (n=49 control; n=17 scrambled shRNA; n=29 IP3R1 shRNA) represented as medians ± interquartile range. A Kruskal-Wallis test was used to assess statistical significance of differences. Control vs IP3R1 KD, p = 2×10^-9^; IP3R1 KD vs scrambled, p = 0.0003. **E**. Scatter plot of max amplitude are represented as medians ± interquartile range. A Kruskal-Wallis test was used to assess statistical significance of differences. Control vs IP3R1 KD, p = 7×10^-9^; IP3R1 KD vs scrambled, p = 3×10^-5^. **F**. FFN511 release was measured in cells stimulated by PACAP. Responses were reported as fusion events per unit area. n=12 cells for scrambled shRNA and n=13 cells for IP3R1 shRNA. Differences between groups were statistically significant (Mann-Whitney, p = 7×10^-7^).

Dispersed cells were subsequently loaded with Cal-520 to monitor changes in cytosolic Ca^2+^ or false fluorescent neurotransmitter (FFN511) to monitor exocytosis (Gubernator et al., 2009). ShRNA-expressing chromaffin cells were visually identified by a TurboRFP reporter. The data show that the Ca^2+^ signals caused by PACAP were substantially smaller in cells with reduced IP3R1 expression (knockdown; KD cells), than in untransfected cells (“control”) or cells with scrambled shRNA. Examples of Cal-520 %ΔF/F records for control, scrambled shRNA, and IP3R1 KD cells stimulated by PACAP are shown in **Figure 9C**. The mean total area under the curve of the PACAP-evoked Ca^2+^ signal was reduced from 8400 ± 2200 in cells with the scrambled shRNA (n = 17) to compared to 1300 ± 160 in KD cells (n = 29) (**Figure 9D**). The mean peak amplitude (± SEM) of the Ca^2+^ increase in KD cells was only 18 ± 2.7% ΔF/F (n = 29) compared to 150 ± 43% ΔF/F in cells expressing the scrambled shRNA (n = 17) (**Figure 9E**).

We also monitored the secretory responses to PACAP in scrambled shRNA and IP3R1 KD groups. As **Figure 9F** shows, reduced IP3R1 expression caused a tenfold reduction in the number of exocytotic events, from approximately 0.05 ± 0.008 events/μm^2^ to 0.005 ± 0.002 events/μm^2^ (p = 7×10^-7^). We also tested whether reduced IP3R1 expression negatively impacts DMPP-evoked Ca^2+^ signals. Interestingly, IP3R1 KD had no appreciable effect on the Ca^2+^ signals caused by DMPP stimulation (**Figures S6A-S6B**).

## Discussion

### Role of Ca^2+^ channels

In this study, we provide strong evidence that Ca^2+^, from two distinct sources – the extracellular space and the ER – is required for PACAP-stimulated secretion to occur (F**igure 10**). Some of the earliest studies to investigate PACAP-stimulated secretion in the chromaffin cell posited a role for extracellular Ca^2+^ in membrane depolarization, fusion, or both (Hill et al., 2011; Mustafa et al., 2007; Przywara et al., 1996; Tanaka et al., 1996). The nature of the pathway by which Ca^2+^ enters cells, however, has remained stubbornly difficult to resolve.

**Figure 10.**
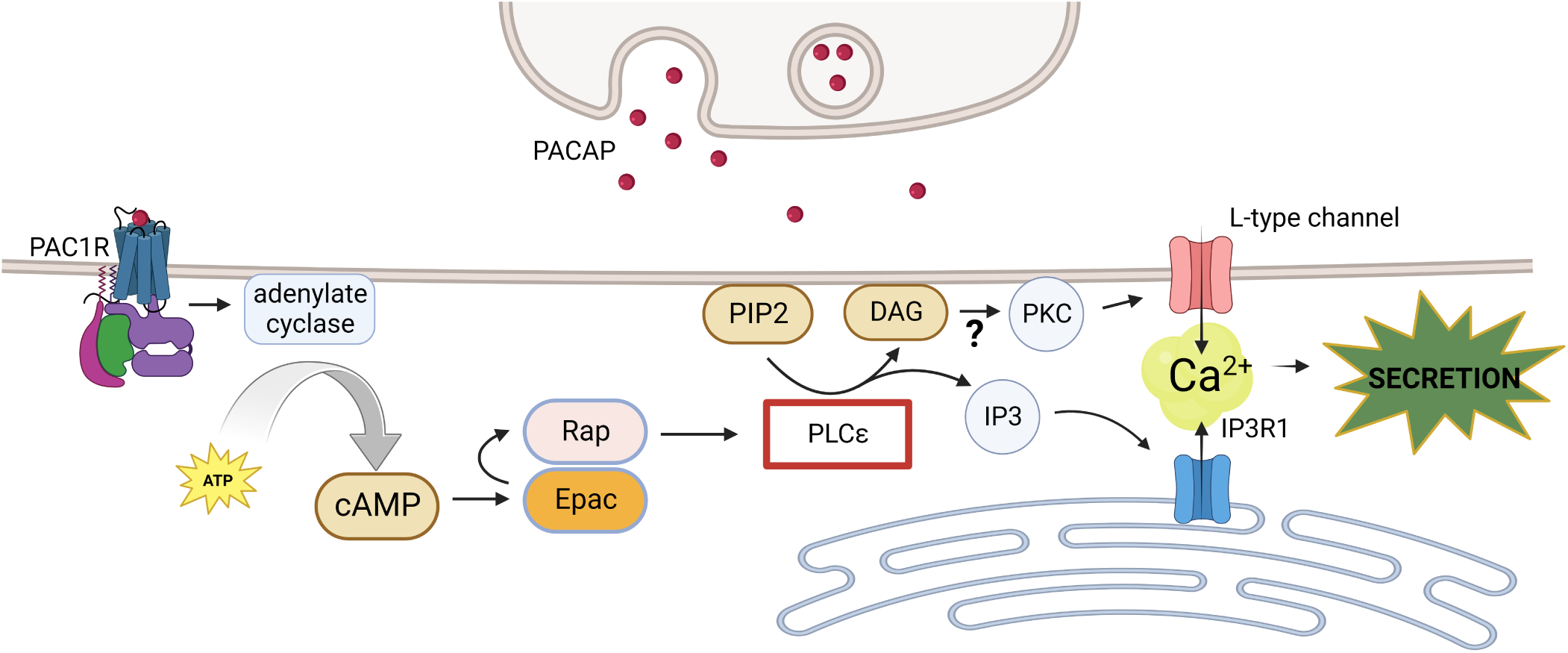
A PACAP-mediated pathway for exocytosis in chromaffin cells. PACAP released from splanchnic terminals onto chromaffin cells causes secretion by harnessing multiple sources of Ca^2+^. PACAP gates the opening of nifedipine-sensitive L-type channels, enabling Ca^2+^ influx into the cytosol, possibly through a mechanism that involves PKC. PACAP also gates Ca^2+^ release from the ER by activating IP3R1s. These pathways depend critically on PLCε activity (boxed). Increases in cytosolic Ca^2+^ are sustained by Ca^2+^-induced Ca^2+^-release (CICR) from the ER. This self-perpetuating CICR causes a persistent secretory response. Cartoon generated with BioRender.

Multiple groups have suggested that Ca^2+^ enters through voltage-gated channels on the plasma membrane of chromaffin cells and PC12 cells (Hill et al., 2011; Osipenko et al., 2000). In fact, it has been proposed that P/Q, N, T, and/or L-type channels are involved in the PACAP pathway (Hill et al., 2011; Osipenko et al., 2000; Tanaka et al., 1996; Taupenot et al., 1999). Although multiple types of voltage-gated channels have been identified in mouse chromaffin cells, those of the L-type are most highly expressed, followed by P/Q and N (Garcia et al., 2006). Here, we assessed the potential contribution of each of these channel families with inhibitors. Only inhibition of L-type channels with nifedipine reduced fluorescent Ca^2+^ signals, PACAP-activated inward current, and exocytosis (**Figures 4**; **Figure S2**). A recent study suggested that T-type channels may play a role in the PACAP secretory pathway (Hill et al., 2011). However, we found no evidence of T-type channel conductances in the mouse chromaffin cell system (not shown). This is not that surprising given the low expression of T-type channels, in general, in the rodent chromaffin cell system (Garcia et al., 2006; Novara et al., 2004).

Through which L-type channel might Ca^2+^ enter the cell? One possibility is the Ca_v_1.3 channel. Chromaffin cells are known to have a relatively depolarized resting potential of around -55 mV (Lingle et al., 2018; Marcantoni et al., 2010; Morales et al., 2023). Channels of the Ca_v_1.3 variety are active at or around the resting membrane potential of the mouse chromaffin cell (Marcantoni et al., 2010). Moreover, chromaffin cells are known to exhibit intrinsic spiking activity in culture, with Ca_v_1.3 regulating the pacemaking current (Marcantoni et al., 2010).

Indeed, cells harvested from Ca_v_1.3 KO mice have substantially reduced spiking activity compared to WT controls (Marcantoni et al., 2010). Interestingly, we have shown that chromaffin cells exposed to PACAP exhibit increased spiking activity (Morales et al., 2023). This suggests a model by which Ca^2+^ influx occurs through nifedipine-sensitive Ca_v_1.3 channels whose activity is upregulated by PACAP.

How might PACAP potentiate Ca^2+^ channels? One possibility is that PACAP slightly shifts their voltage-dependence to more negative potentials, thereby increasing their open probability at or near the resting membrane potential. Another possibility is that PACAP activates a separate conductance to depolarize the membrane potential and activate Ca_v_s in that way (Inoue et al., 2020; Kuri et al., 2009). Both possibilities may involve activation of PKC downstream of DAG (Smrcka et al., 2012) (**Figure 10**). Although a role for PKC was not investigated here, it was previously shown to activate depolarizing currents in chromaffin cells (Kuri et al., 2009).

### The role of ER Ca^2+^

PACAP-stimulated Ca^2+^ signals fluctuate with a variable amplitude and time course. Because these Ca^2+^ signals are also long-lived (i.e., do not desensitize during PACAP exposure), we speculated that they were unlikely to be generated solely by voltage-gated Ca^2+^ channel activity. That internal Ca^2+^ stores could be involved in the PACAP pathway has been suggested by others (Mustafa et al., 2010; Payet et al., 2003; Tanaka et al., 1996). Experiments in bovine chromaffin cells, rat chromaffin cells, and even human fetal chromaffin cells have shown PACAP to harness internal Ca^2+^ to regulate release (Mustafa et al., 2010; Payet et al., 2003; Tanaka et al., 1996). However, most previous studies have focused on the potential role of RyRs, but not IP3Rs (Mustafa et al., 2007; Payet et al., 2003; Tanaka et al., 1996). In our studies, inhibition of RyR function had no effect on the properties of the Ca^2+^ signals stimulated by PACAP (**Figures 5A-5B**). Moreover, only transcripts for RyR2 and RyR3 were detected in mouse chromaffin cells. RyR transcript expression was also much lower than that of the IP3R1 isoform (**Figure 7**).

Whereas inhibition of RyRs had no effect on PACAP-stimulated Ca^2+^ signals, IP3R inhibition caused substantial reductions in their magnitude (i.e., area under the curve) and maximum amplitude (**Figures 5A-5B**). Genetic depletion of the most highly expressed IP3R isoform in chromaffin cells – IP3R1 – had profound detrimental effects on Ca^2+^ signals and exocytosis (**Figure 9**). The involvement of IP3Rs is consistent with known activities of PACAP in chromaffin cells. PACAP-stimulated secretion requires intact signaling through PLCε (Chen et al., 2023; Morales et al., 2023). In addition, PLCε has been shown to cause DAG and IP3 generation in other systems (Citro et al., 2007; Lucchesi et al., 2016; Smrcka et al., 2012). This suggests that a key function of PLCε in chromaffin cells is to generate IP3 and activate the IP3 receptor.

### The relationship between Ca^2+^ influx and ER Ca^2+^ release

Our results reveal a close functional relationship between extracellular Ca^2+^ levels and the Ca^2+^ stored in the ER. Removal of Ca^2+^ from the external medium caused an almost immediate decrease in ER Ca^2+^ levels (**Figures 3C-3D**). One interpretation of these results is that the nifedipine-sensitive Ca^2+^ channels controlling Ca^2+^ influx are organized to rapidly signal their status to the portions of the ER abutting the plasma membrane. Thus, Ca^2+^ coming into the cell might serve two purposes: 1) to stimulate Ca^2+^ release from the ER; and, 2) to provide a source of Ca^2+^ through which ER stores are replenished. Such an organization is reminiscent of the relationship between Ca^2+^ entry sites on the plasma membrane and RyRs on the sarcoplasmic reticulum (Scoote, 2002). A spatial coupling between Ca^2+^ entry sites and ER release sites is economical from the point of view of the cell. A relatively small Ca^2+^ influx of the sort stimulated by PACAP (**Figure S2A**) is great amplified due to the action of Ca^2+^ in sensitizing Ca^2+^ release from IP3Rs (i.e., CICR) (Foskett et al., 2007; Lock et al., 2018; Lock and Parker, 2020; Smith and Parker, 2009; Smith et al., 2009). This results in a large, prolonged Ca^2+^ signal that provokes a correspondingly long-lived secretory response in the chromaffin cell (**Figure 10**). Support for such a model is substantiated by confocal imaging of the ER network (i.e., with ER-GCaMP6-150) that shows it be extensively branched with varicosities rich in Ca^2+^ appearing at or near the plasma membrane (**Figures 3A-3B**). IP3R1 is frequently co-localized with an ER marker, KDEL, even in regions of the ER close to the plasma membrane (**Figure 8**).

### Distinct roles of PACAP and ACh in the context of the sympathetic stress response

ACh-based stimulation of chromaffin cells causes a burst of fusion events that decrease in frequency over time (Morales et al., 2023). Secretion stimulated by PACAP has a distinct kinetic profile; fusion events occur at a regular frequency without evident rundown or desensitization (Morales et al., 2023). In addition to phenomenological differences in the secretory response, ACh and PACAP also rely on different signaling pathways to cause secretion. For example, we previously showed that the absence of PLCε has little to no effect on the secretory response stimulated by ACh (which activates nicotinic and muscarinic receptors), DMPP (a nicotinic agonist), or bethanechol (a muscarinic agonist) (Chen et al., 2023).

Meanwhile, the absence of PLCε effectively eliminates secretion triggered by PACAP (Chen et al., 2023; Morales et al., 2023). Here, our previous results were expanded to show that disruption of IP3R signaling inhibits secretion triggered by PACAP, but not DMPP. Note, only the effect of DMPP stimulation was evaluated here as most ACh-stimulated release, *in situ*, occurs via activation of nicotinic receptors (Wakade and Wakade, 1983). Based on these results, we conclude that the two major pathways for secretion in the adrenal medulla – one stimulated by ACh and the other stimulated by PACAP – have little functional overlap. Of course, what we are measuring here are responses to acute stimulation. Longer-term stimulation of chromaffin cells in culture, or the adrenal medulla, *in vivo*, might uncover novel forms of crosstalk between the pathways. Nevertheless, the implications of our results for the secretory response are profound. They suggest secretion from the adrenal medulla is under the control of multiple neurotransmitters signaling through independent signaling pathways. Such a system may enable, for example, a range of secretory responses from chromaffin cells to stressors that differentially activate the sympathetic nervous system.

## Acknowledgements

This study was supported by R01NS122534 (AA), R01MH125849 and R21NS126779 (PJK), R01GM136826 (RGM), R35GM127303 (AVS), DE019245 (DIY). BLC is supported by the NIH G-RISE program, T32GM144873. The authors have no conflicts to report.

## Supplementary Figure Legends

**Figure S1.**
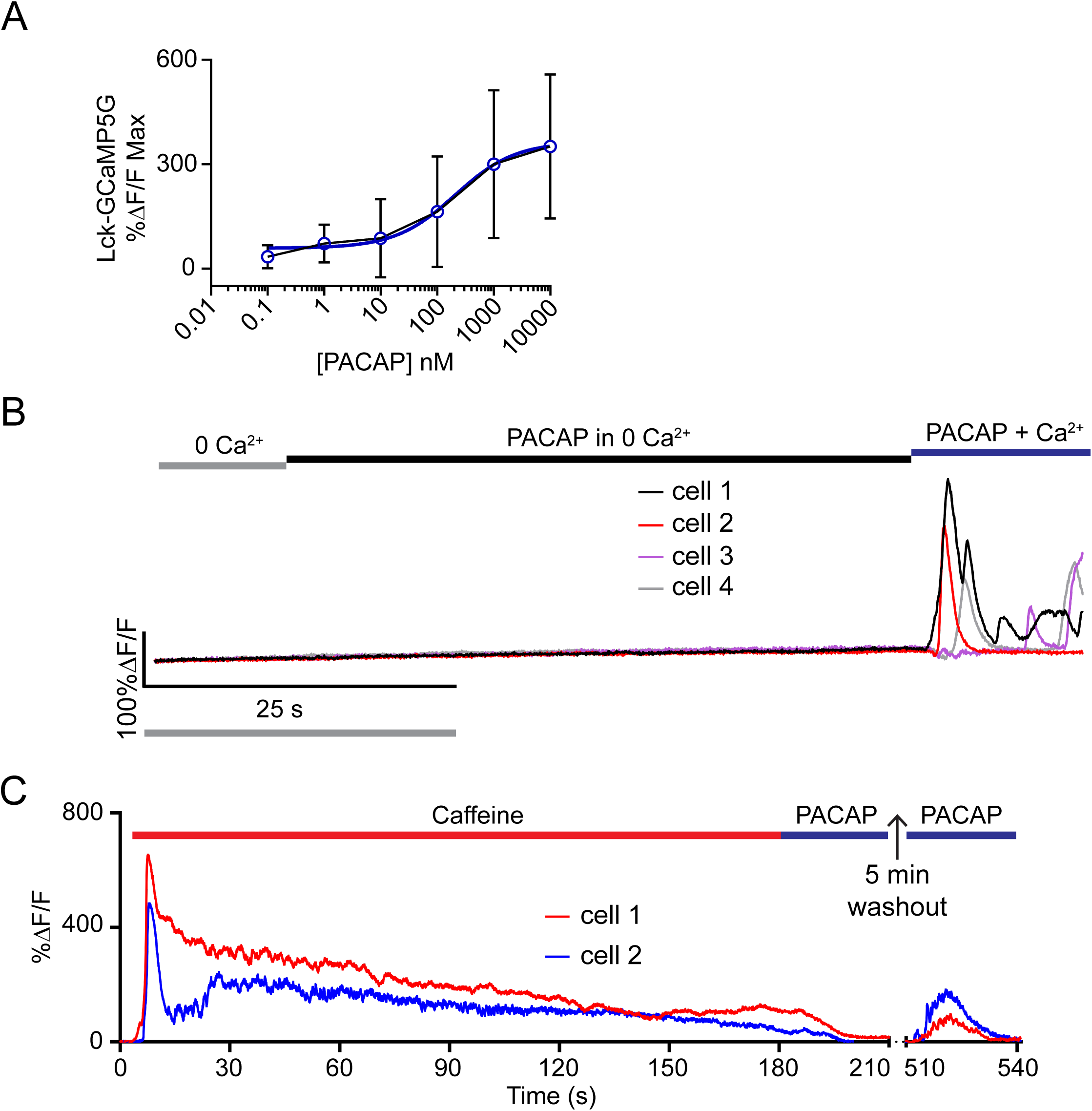
Examples of PACAP-stimulated Ca^2+^ transients. **A**. PACAP-stimulated dose-dependent increases in Lck-GCaMP5G fluorescence in chromaffin cells (0.1 nM to 10,000 nM). Data points show means ± SD. Data were fit with an [agonist] vs response curve using GraphPad Prism. **B**. Additional examples of %ΔF/F versus time records showing external Ca^2+^ is required for PACAP Ca^2+^ transients. **C**. Two additional representative traces showing that ER Ca^2+^ depletion with caffeine inhibits PACAP-stimulated Ca^2+^ transients.

**Figure S2.**
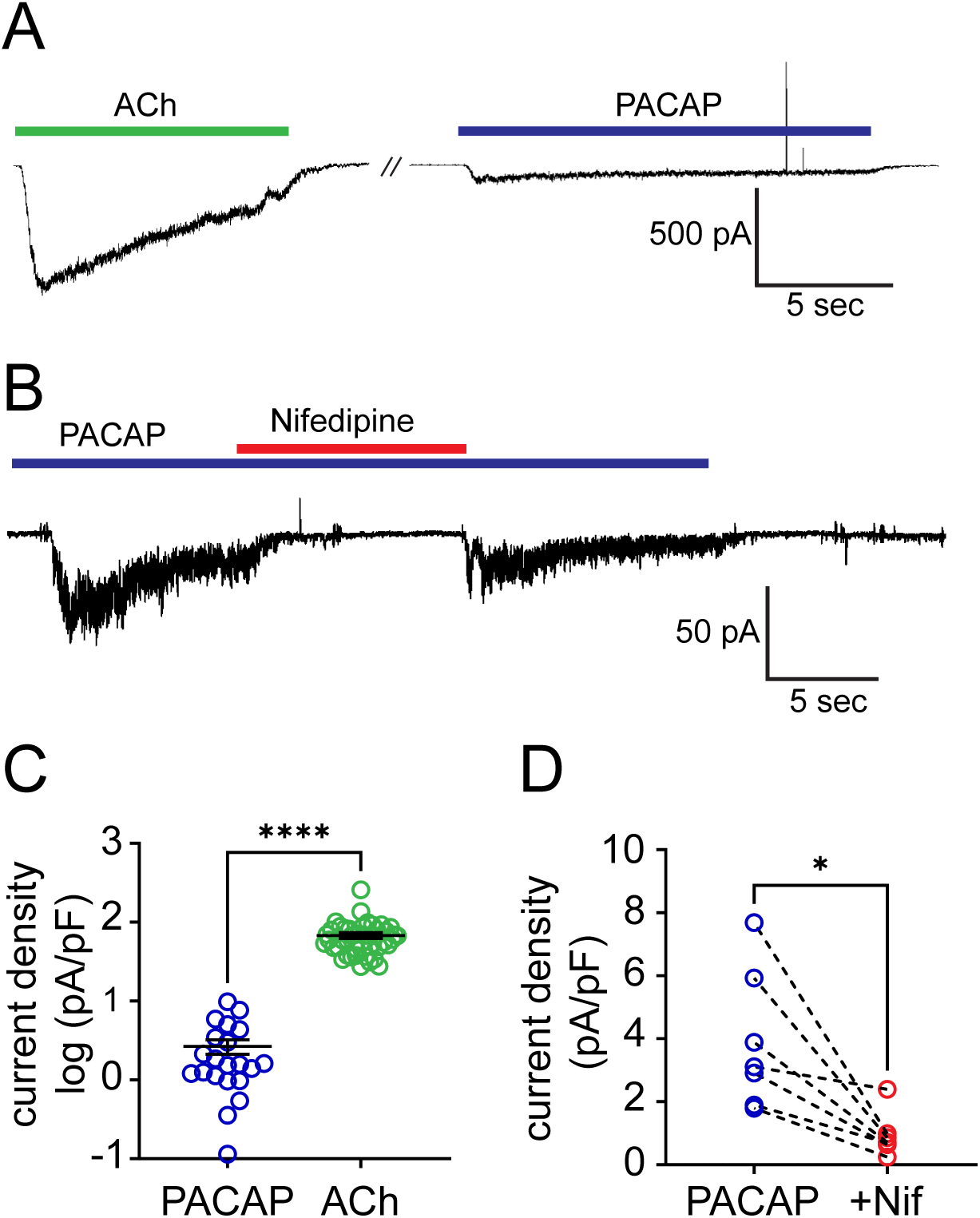
PACAP-mediated currents were nifedipine sensitive. **A**. PACAP (500 nM) activated a slowly desensitizing current in mouse chromaffin cells (n=12) whereas application of acetylcholine (100 nM) evoked a rapidly desensitizing current in all cells (n=44). Cells were maintained at 35°C and held at a membrane potential of -55 mV. **B**. Co-application of nifedipine (10 μM) was shown to diminish the current to baseline levels. When nifedipine was removed, the PACAP-evoked current partially recovered (n=7). **C**. Averaged peak amplitudes of acetylcholine- and PACAP-evoked currents normalized to whole cell capacitances (pA/pF). PACAP-mediated currents had an average peak amplitude of 2.7 pA/pF whereas acetylcholine-mediated currents had an average peak amplitude of 68 pA/pF. **D**. Application of nifedipine significantly reduced PACAP-mediated currents from an average of peak value of 3.9 pA/pF to 0.93 pA/pF. Statistical significance (student’s t-test) indicated by asterisk: (****) p-value = <0.0001. (*) p-value = 0.004.

**Figure S3.**
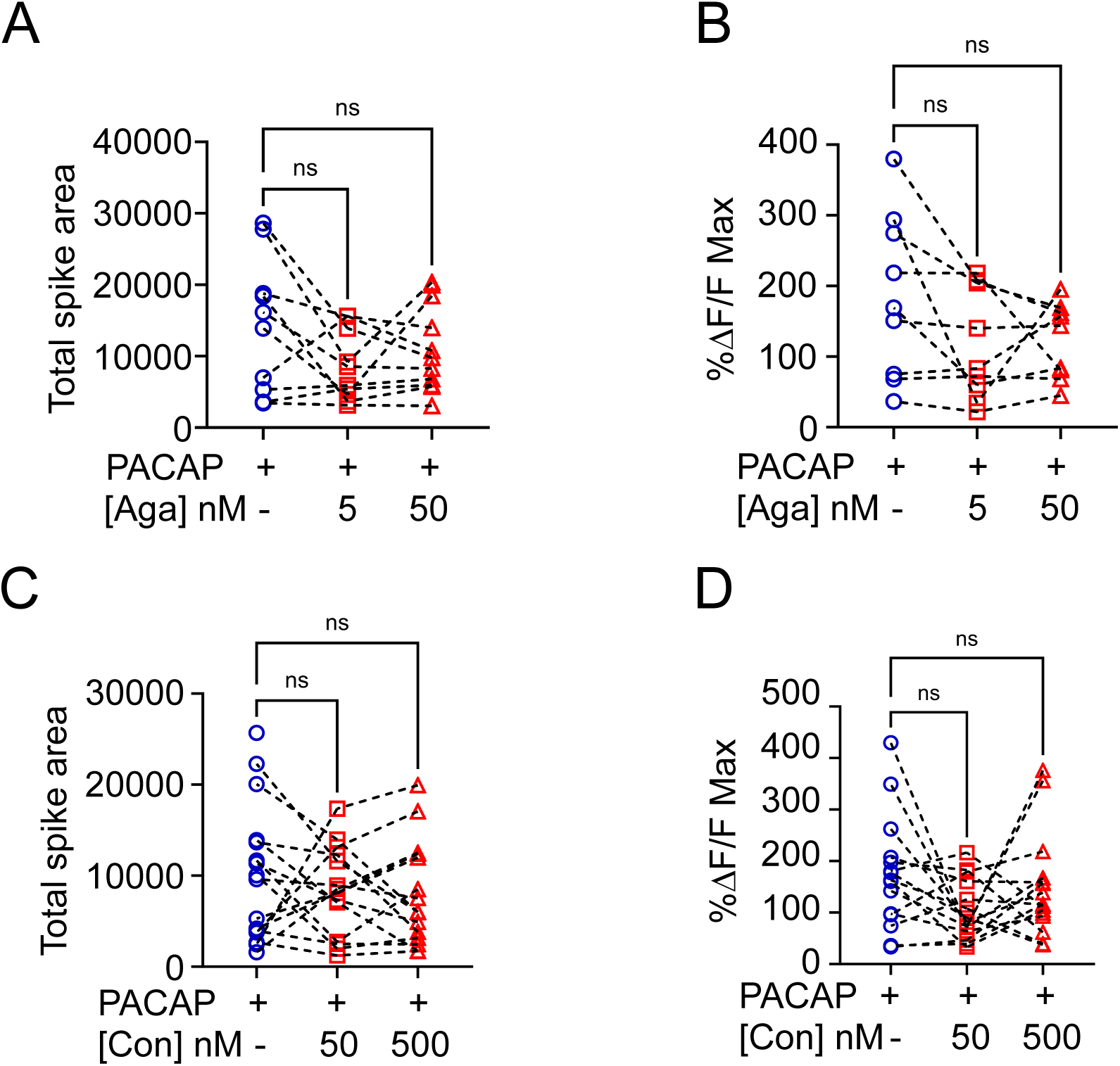
P/Q-type and N-type channel blockers did not inhibit PACAP-evoked Ca^2+^ signals. **A** and **B**. The total Ca^2+^ spike area and maximum %ΔF/F for chromaffin cells stimulated with 500 nM PACAP alone or with two different ω-Agatoxin (P/Q-type channel blocker) concentrations; n=9. Differences between groups were not significantly different (Kruskal-Wallis; p > 0.05). **C** and **D**. The total Ca^2+^ spike area and maximum %ΔF/F for chromaffin cells stimulated with 500 nM PACAP alone or with two different ω-Conotoxin GVIA (N-type channel blocker) concentrations; n=15. Differences between groups were not significantly different (Kruskal-Wallis; p > 0.05).

**Figure S4.**
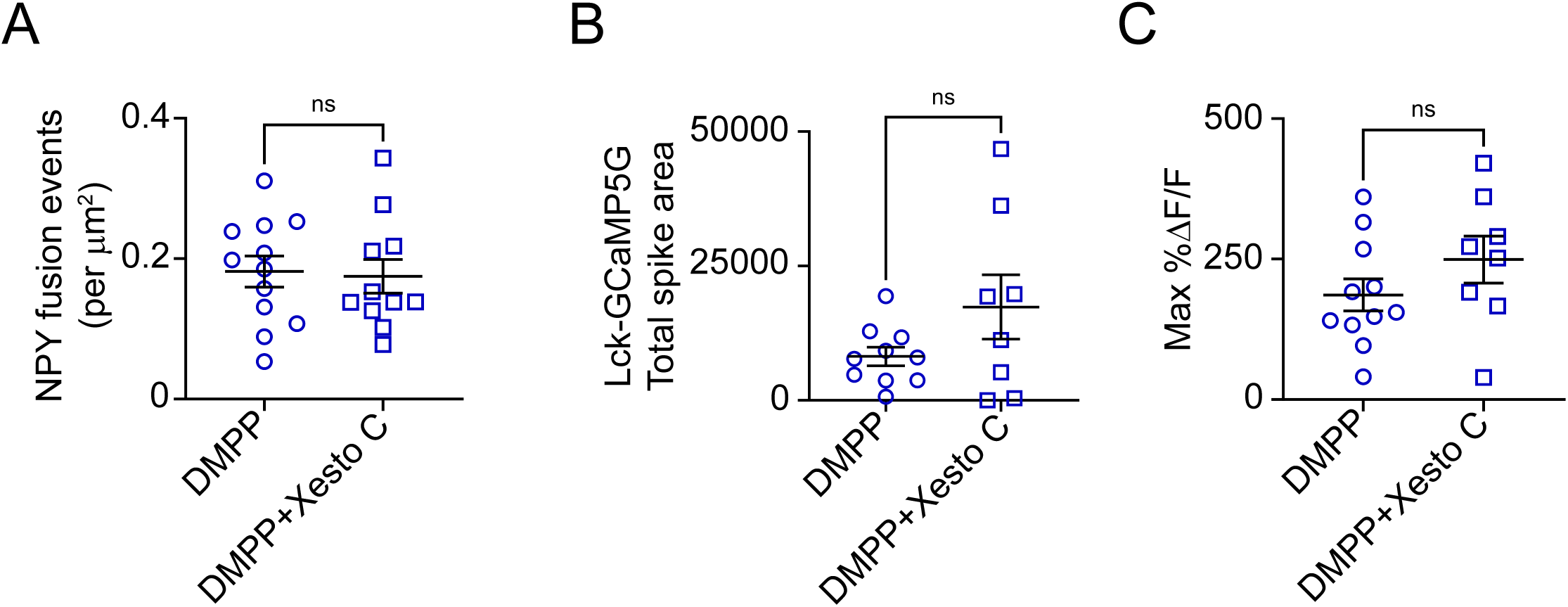
Xestospongin C did not inhibit DMPP-evoked Ca^2+^ transients or secretion. **A**. Cells were perfused with either 1 μM DMPP alone (n=12) or DMPP+Xesto C (5 μM; n=11), after a 5 min incubation with Xestospongin. NPY fusion events per unit area were not different between groups (Student’s t-test; p > 0.05). **B** and **C.** Neither the total Ca^2+^ spike area nor maximum %ΔF/F for chromaffin cells were different between groups (Student’s t-test; p > 0.05). n=11 for DMPP, n=8 for DMPP+Xesto C.

**Figure S5.**
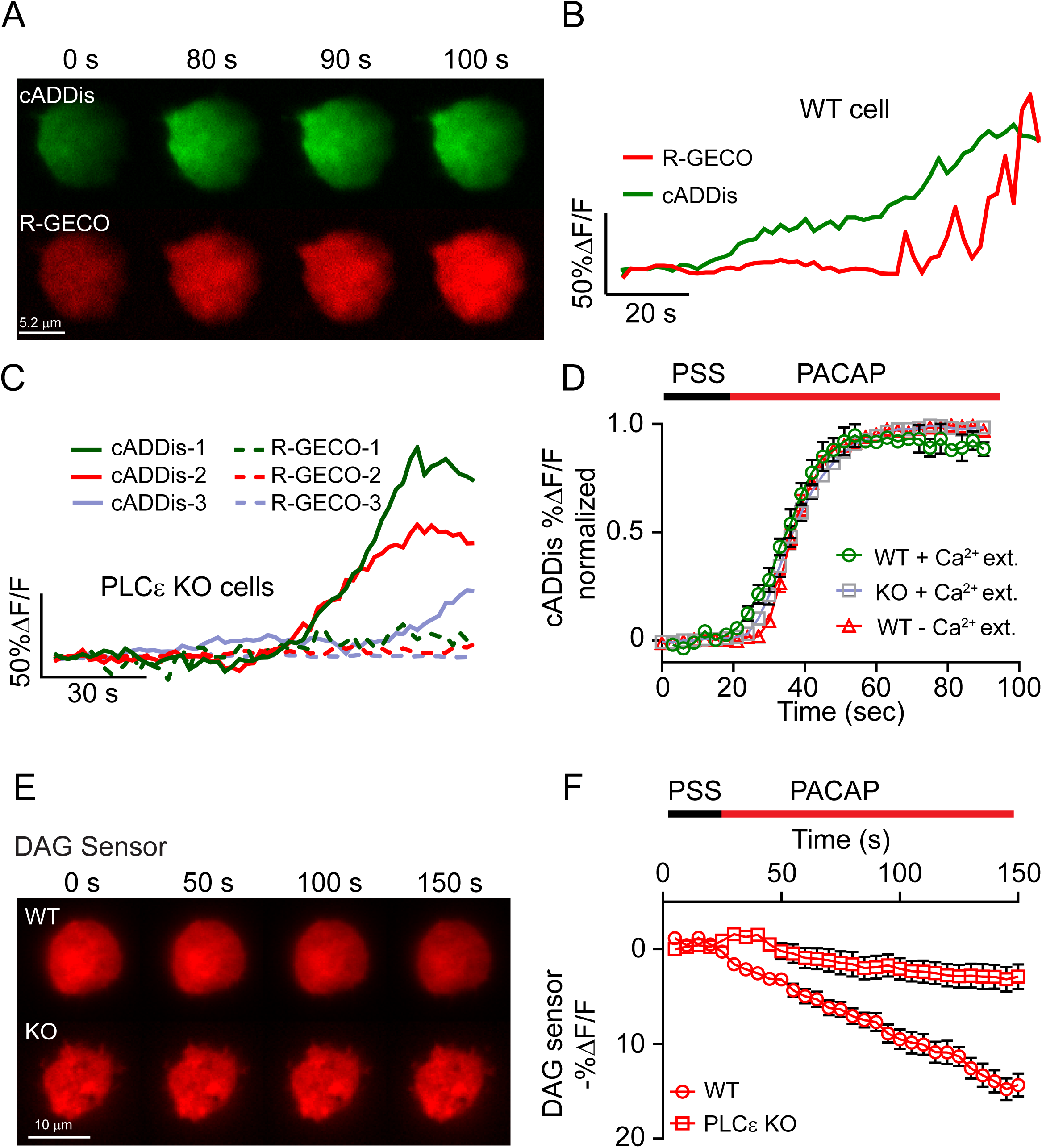
cAMP was produced in PLCε KO cells but not DAG or Ca^2+^ signals. **A.** PACAP stimulation caused increases in cADDis (cAMP indicator) and R-GECO (Ca^2+^ indicator) fluorescence (n = 5). **B**. cADDis and R-GECO intensity versus time records from cell in **A**. **C**. Three representative records showing that PLCε KO inhibited PACAP-evoked Ca^2+^ transients but not cAMP production (n=5). **D**. cAMP production (measured via cADDiS fluorescence) is normal in PLCε KO cells stimulated by PACAP with or without extracellular Ca^2+^. The three curves are essentially overlapping (student’s t-test; p > 0.05). Individual data points on curves represented as means ± SEM. n=13 for PACAP+Ca^2+^ stimulation of WT cells; n=24 for PACAP+Ca^2+^ stimulation of PLCε KO cells; n=11 for PACAP-Ca^2+^ stimulation of WT cells. **E**. Examples of WT (n=26) and KO (n=26) cells expressing a downward red DAG sensor (Montana Molecular) stimulated by PACAP (n=26). F. DAG fluorescence decreased in WT cells after PACAP application, but not in KO cells. Differences between data sets were statistically significant (p < 0.05) after 30 s.

**Figure S6.**
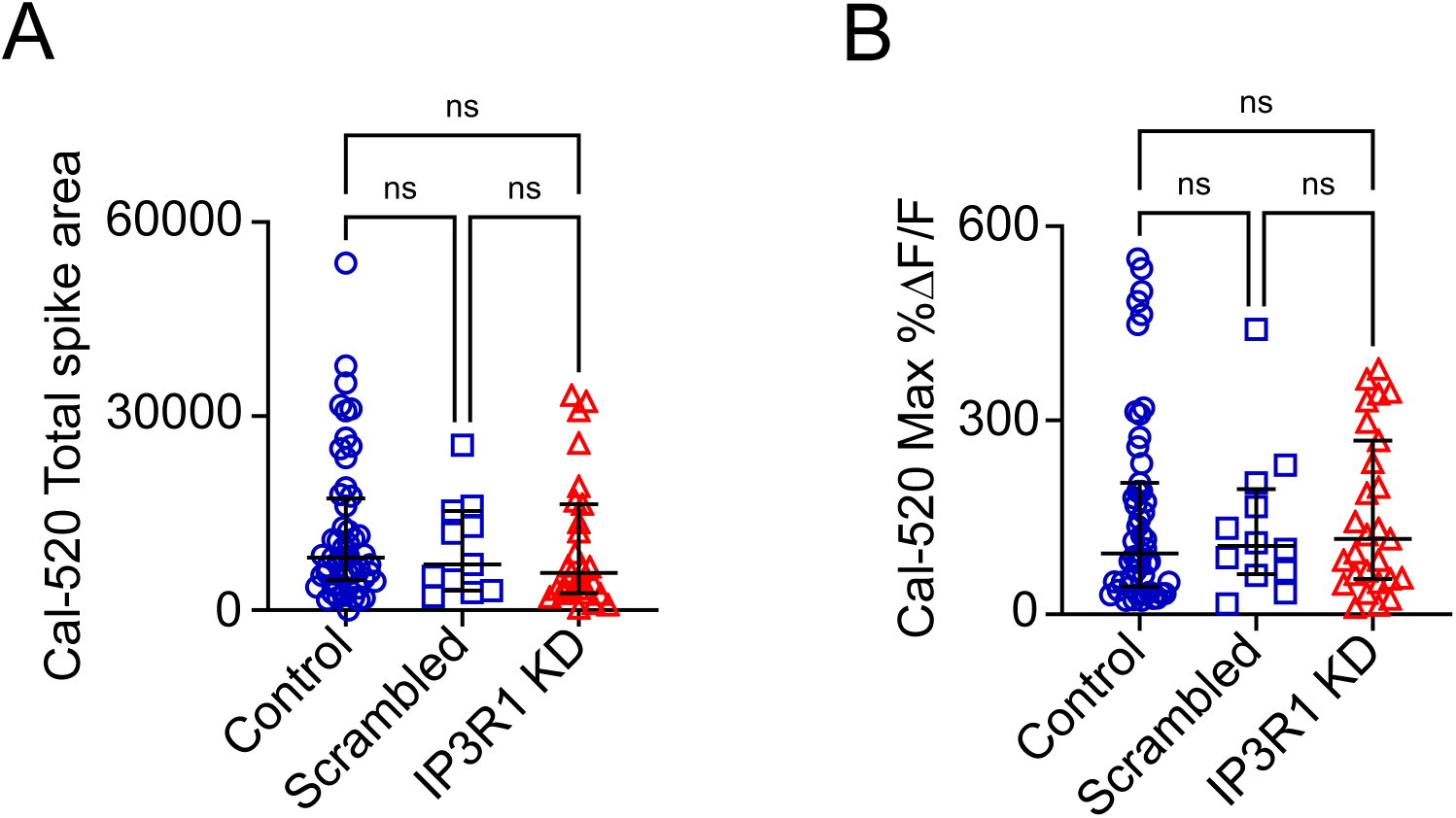
DMPP-evoked Ca^2+^ transients were not reduced in IP3R1 KD cells. **A** and **B.** Scatter plot of total spike area and maximum %ΔF/F of Cal520 fluorescence. Cells were stimulated by 1 μM DMPP. n=51 control (untransfected), n=12 for scrambled shRNA, n=27 for IP3R1 KD. Data are shown as medians ± interquartile range. Differences between groups were not statistically significant (Kruskal-Wallis; p > 0.05).

## Notes

### Competing Interest Statement

The authors have declared no competing interest.

